# Hemispheric lateralization in older adults who habitually play darts: a cross-sectional study using functional near-infrared spectroscopy

**DOI:** 10.1101/2022.03.18.484828

**Authors:** Koki Toyofuku, Satoru Hiwa, Kensuke Tanioka, Tomoyuki Hiroyasu, Masaki Takeda

**Author notes:** **Correspondence:** Satoru Hiwa.

## Abstract

Exercise training integrating physical and cognitive activities is gaining attention because of its potential benefits for brain health. This study focuses on exercise training using a dart game called Wellness Darts. Wellness Darts is a sport involving throwing darts and walking to pull them out of the board, memorizing the score, and subtracting it from the total score, thus allowing the performance of two tasks simultaneously: exercise and calculation. It is expected to maintain and improve cognitive function, and whether the continual dart training affects brain function is of great interest. Before conducting the longitudinal study revealing its effect on brain function, we aimed to cross-sectionally confirm the difference in hemispheric lateralization between expert and non-expert players. Functional near-infrared spectroscopy (fNIRS) was used to measure brain activity for three groups: an expert older group who practiced darts continually, a non-expert older control group, and a non-expert younger control group. Their brain activity patterns were quantified by lateralization index (LI) and compared between groups. The results showed that the younger and the expert older groups had significantly higher LI values than the non-expert older group and did not differ between the expert older and the younger groups. Our results suggest that the Wellness Darts possibly promote hemispheric lateralization.

## 1 Introduction

Cognitive decline is one of the most crucial issues to be addressed in the aging population. Various interventions have been proposed to maintain and improve cognitive function in older adults, including physical activities and cognitive exercises. In this context, exercise and sports that integrate physical and cognitive activities are gaining attention because of their potential benefits for brain health.

Wu et al. found that Mind-Body exercise, particularly tai chi and dance, improves global cognition, cognitive flexibility, working memory, verbal fluency, and learning in cognitively impaired and non-disabled older adults [1]. Furthermore, Babaei et al. suggested that frequent moderate aerobic activity can be associated with improved neurocognitive performance for older adults [2]. Gheysen et al. stated that combined physical and cognitive activity programs should be promoted to prevent and treat cognitive decline in older adults [3]. Similarly, Rieker et al. found that performing cognitive and physical exercise simultaneously and interactive training produced the most significant gains in executive functions, speed, and global cognition, and the most significant improvements in physical functions [4]. Also, physical activity associated with exercise training has been shown to cause substantial changes in brain health and cognitive performance, potentially affecting a wide range of cognitive functions, including memory, attention, and executive function [5]. Additionally, Liu et al. suggested exercise-induced enhancement of executive function in young individuals, providing practical evidence to encourage regular exercise to maintain cognitive function and promote brain health [6]. These previous studies imply that exercise training, especially the combined physical and cognitive activity program, is one of the most effective interventions for brain health.

This study focused on exercise training using a darts game called Wellness Darts. Darts is one of the sports that seek fun and gameplay by subtracting the score obtained by the player from the points they have and reducing it to zero. Wellness Darts also has the unique feature of calculating scores by rote [7]. “Rote,” in this case, refers to the process of quickly mentally calculating the score of a dart thrown by a player on a dartboard and subtracting it from the points the player has. Wellness Darts is a combined physical and cognitive activity program that involves not only throwing darts but also walking to pull them out of the board, memorizing the score, and subtracting it from the total score, thus allowing the performance of two tasks at the same time: exercise and calculation [7]. Simultaneous performance of the motor and cognitive tasks, each effective in improving cognitive function as a single task, is more effective in increasing PFC activity and improving cognitive function than performing either task alone [8]. Takeda et al. conducted cognitive tests before, one month, and two months after an experiment on a Wellness Darts intervention group and a control group of older adults [7]. The results showed that continuing Wellness Darts increased older adults’ short-term memory test (STM test) scores. However, the impact of continual practice of Wellness Darts on brain function is still being determined. To develop Wellness darts as a combined physical and cognitive activity program, it is essential to understand their potential effects on brain function.

This study investigates the cognitive status and brain function of participants who practiced Wellness Darts in older adults. Their characteristics of brain function and the performance of cognitive tests will be compared cross-sectionally among older adults who are proficient with darts, older adults who are inexperienced with darts, and younger adults who are inexperienced with darts. The Hemispheric Asymmetry Reduction in OLDer adults (HAROLD) model proposed by Cabeza et al. asserts that the degree of Frontal lobe (FL) lateralization seen in youth decreases with age [9, 10, 11,12, 13, 14, 15, 16]. In addition, Erickson et al. reported that when cognitive training was conducted on healthy older adults, lateralization was observed with improved cognitive performance, bringing the brain activation pattern closer to that of younger adults [12]. That is why increased lateralization can be used as a measure of cognitive improvement. On the other hand, brain regions such as the inferior frontal gyrus (IFG), superior frontal gyrus (SFG), middle frontal gyrus (MFG) [17], and DLPFC have been reported to be activated during mental calculation [18]. Furthermore, the inferior parietal lobule (IPL) in patients with cognitive decline, such as MCI and Alzheimer’s disease, is reported to be less activated [19].

Based on these findings, we proposed the following three hypotheses: (1) older adults who practice Wellness Darts (referred to as the ‘expert older group’) will show a level of hemispheric lateralization similar to that of younger people who had never played Wellness Darts before (referred to as ‘younger group’ as a control), with activation of the dorsal superior frontal gyrus (SFGdor) and MFG, which are known to be activated during mental arithmetic. (2) There is no difference in the level of IPL activation between the expert older group and the younger group. (3) There is a positive correlation between the level of hemispheric lateralization in brain activity and the number of days of Wellness Dart experience in the expert older group. These three hypotheses were tested by measuring brain activity using functional near-infrared spectroscopy (fNIRS), a non-invasive functional brain imaging system robust to body movement. The third group, the older adults who had never played Wellness Darts (referred to as the ‘non-expert older group’), was used as the older control to confirm that hypotheses (1) and (2) were valid only for the expert older group.

## 2 Materials and Methods

### 2.1 Participants

Twenty-one healthy younger adults (13 males, seven females; mean age: 22.5 ± 0.8 years) who had never played Wellness Darts, 21 healthy older adults (8 males, 13 females; mean age: 75.1 ± 4.3 years) who continually played Wellness Darts, and 21 healthy older adults (4 males, 17 females; mean age: 74.1 ± 4.5 years) who had never played Wellness Darts participated in the experiment. These three participant groups were referred to as the younger group, the expert older group, and the non-expert older group. The participants for the expert older group were recruited from Wellness Darts players in the Kyotanabe-Doshisha Sports Club (KDSC), one of the community sporting clubs in Kyoto, Japan. The participants for the non-expert group were also recruited based on the acquaintanceship of the members of the expert older group. Only healthy participants over 60 could participate in this study for these two groups. For the younger group, the healthy undergraduate and graduate school students aged over 20 were recruited from Doshisha University, Kyoto, Japan. The sample size for each group was determined with a significance level of 0.05 and a power of 0.8 to detect the effect size of Cohen’s *d* = 0.79 reported in a previous study comparing the degree of hemispheric lateralization in older adults using fNIRS [**Error! Reference source not found.**] (G*Power was used for the calculation).

### 2.2 Wellness Darts task

Wellness Darts is a darts game developed by the Japan Wellness Darts Association. The standard game of Wellness Darts is the “Zero One Game 251”. In the game, each player is assigned a score of 251 and throws darts until the score reaches zero, subtracting the score of each dart thrown from the holding points. The first player to get a score of exactly 0 wins. Each player can throw darts three times in their turn (which is defined as one set) and is required to make their score exactly zero on the third throw for either of five sets. In addition, each player must subtract the points earned from their score by themselves. We assumed that these mental arithmetic operations, the planning of throws to reach zero as quickly as possible, and the exercise associated with throwing darts could be effective in improving cognitive function because the Simultaneous performance of the motor and cognitive tasks said to be effective in increasing PFC activity and improving cognitive function [8]. However, since the official rules of Wellness Darts place a physical burden on the participants due to the long duration of the experiment, in this study, the initial score of the subjects was set at 128 points, and each subject played the game for a maximum of five sets. Furthermore, the game was terminated even if the score did not reach zero in the fifth set.

### 2.3 Procedure

Participants played up to five sets of Wellness Darts with a maximum of 128 holding points while brain activity and head acceleration were simultaneously recorded. As described below, the accelerations were used to remove noise due to head movements. In each of the three throws in one set of the Wellness Darts game, the participants were instructed by the recorded voice prompts and the beep cue from the experimental computer on when to hold the dart, throw, and start the calculation; the participants performed each action following the instructions.

During the planning block set at the beginning of each throwing session, participants were asked to determine where to hit (designate scoring points) within 5 s for the first throw and within 13 s for the remaining two throws. At the beginning of the second and third throwing sessions, they were provided an extra 8 s in the planning blocks to tally the points from the previous throws. Of note, we excluded the beginning 8 s. Instead, we applied the later 5 s for the subsequent analyses because we assumed that the first throw did not involve the confirmation of the hit point and employed different cognitive processes than those used in determining where to hit. The planning block was followed by dart-throwing within 3 s after the beep cue.

Following dart throwing, the participants were instructed to move their gaze away from the dartboard to the fixation point and not to think of anything in particular. This waiting block consisted of a random duration between 28 - 32 s to reduce the anticipatory response. After completing the three throwing sessions, the participants commenced the calculation block, where they were instructed to subtract the scores, they received during the previous throws from the total score and write the remaining points on the scoring sheet. Then, they were asked to press a button to proceed to the 20 s rest block. During the rest block, they stayed the same as in the waiting block. This procedure was repeated for five sets. The overall procedure is illustrated in Figure 1.

**Figure 1.**
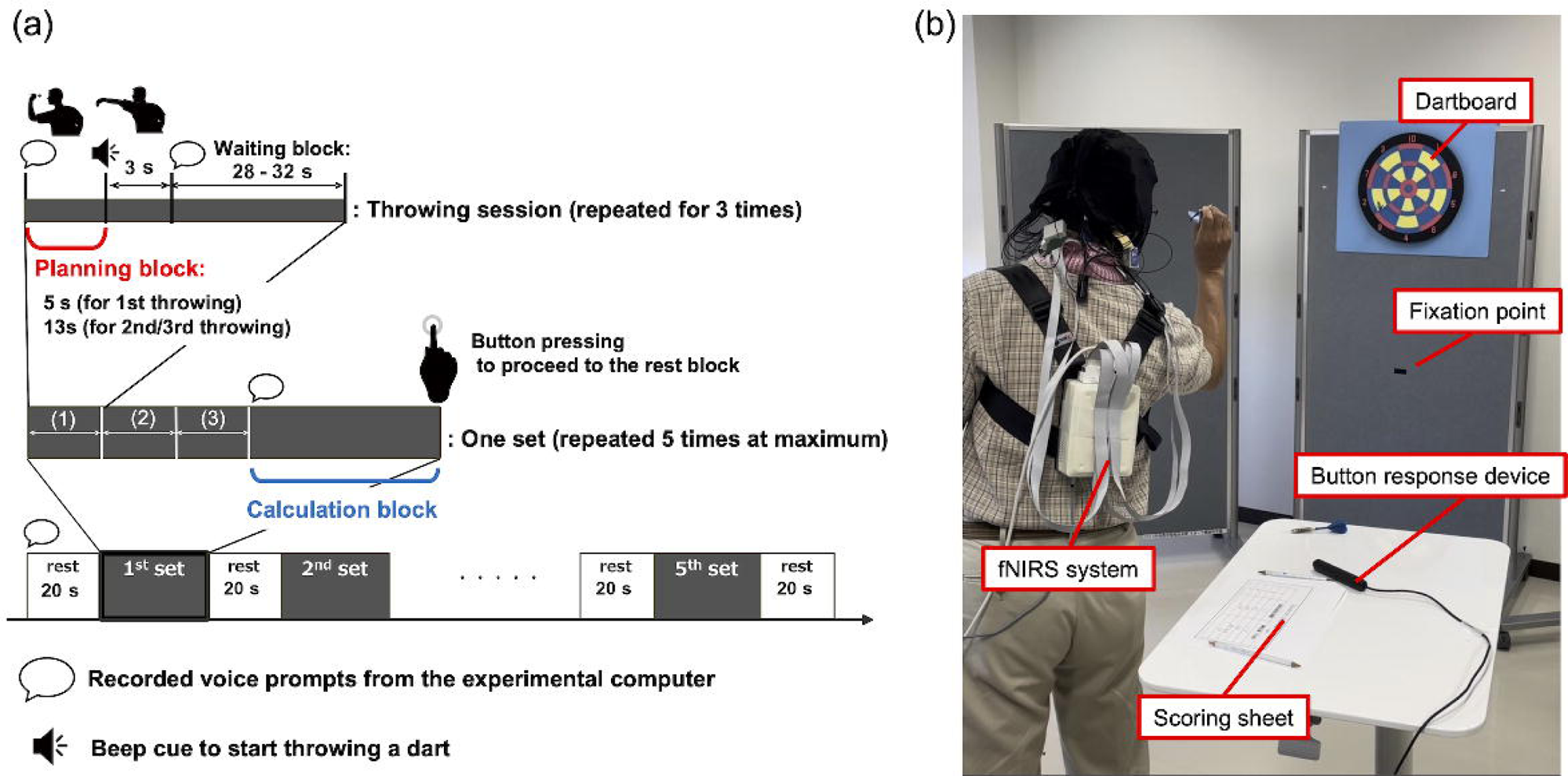
Overview of the experiment. A. The experimental procedure. B. The experimental setup.

All participants were informed of the methods and risks of the experiment and signed a written informed consent form. This study was conducted in accordance with the Research Ethics Committee of Doshisha University (Kyoto, Japan) (approval code: 20014). Before the experiment, participants were asked to complete a questionnaire, a dart-throwing performance test, and five cognitive tests: Simple Cognitive (SC) test, Computational Ability Test, Short-Term Memory (STM) Test, Trail Making Peg Test (TMPT), and CogEvo. Participants then performed one set of dart-throwing before being fitted with the fNIRS device to understand the experimental procedure.

### 2.4 fNIRS data acquisition

Changes in the concentration of oxyhemoglobin (HbO), deoxyhemoglobin (HbR), and total hemoglobin (HbT) in the participants were measured using NIRSport2 (NIRx Medical Technologies, LLC). The sampling frequency was 4.36 Hz, and the wavelengths of the near-infrared light were 760 and 850 nm, respectively [21]. As shown in Figure 2 (b), 16 sources and 14 detectors were placed on the frontal, parietal, and temporal regions of the head. A pair of sources and detectors adjacent at 30-mm intervals constituted one standard channel, and 44 standard channels were placed. In addition, a short-distance detector bundle (NIRx Medical Technologies, LLC) was established to eliminate extra-cerebral blood flow alterations [22, 23]. The signals obtained from the 16 short-distance channels were used to detect extracerebral signals and to remove the effect of skin blood flow in the signals measured by the standard channels. To remove noise from body motion, a six-axis accelerometer (NIRx Medical Technologies, LLC) was used to measure head acceleration at a sampling frequency of 4.36 Hz, the same frequency as the change in hemoglobin concentration [24]. The accelerometer was placed at the top of the head, as shown in Figure 2 (b). Presentation software (Neurobehavioral System Inc.) was used to control the presentation and timing of the recorded voice prompts and the beep cue during the experiment, and the signal measurement and execution of the experimental task were synchronized. The 3D coordinates of the light source and detection probe were measured using a 3D magnetic space digitizer (Fastrak, Polhemus, USA) to estimate the exact measurement position of the brain area for each individual. The measured 3D coordinates were entered into the ‘register2polhemus’ function, and the ‘depthmap’ function implemented in the NIRS toolbox [25], and the distance from each standard channel to the brain surface was calculated. On the basis of this distance, the regions of the brain defined by the automated anatomical labeling atlas [26] corresponding to each standard channel were identified.

**Figure 2.**
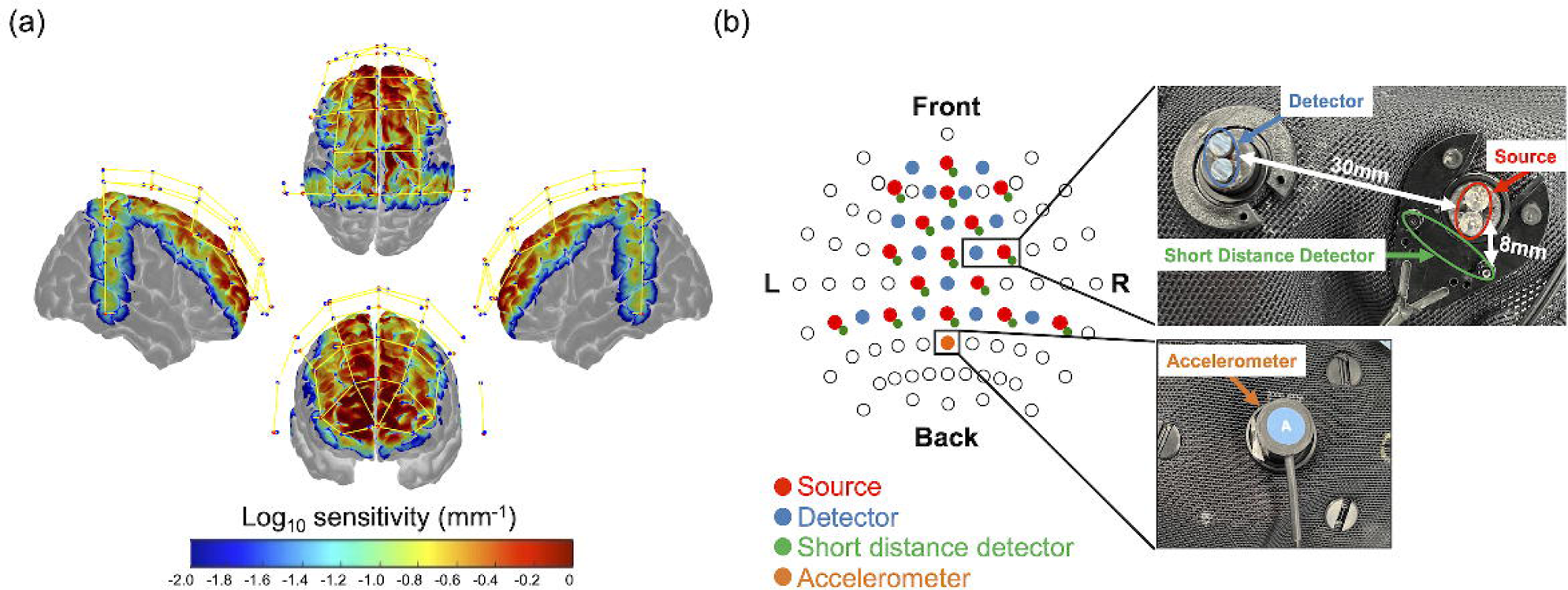
Probe displacement of fNIRS. (a) Monte-Carlo simulation results over the frontal cortex, temporal cortex, and parietal cortex. Red dots represent emitters, blue dots represent detectors, and yellow lines represent measurement channels, respectively (created using Homer 2 AtlasViewer; v2.8, p2.1: https://www.nitrc.org/frs/shownotes.php?release_id=3956). The color bar represents the spatial sensitivity of fNIRS measurements. (b) The two-dimensional fNIRS montage using the International 10-20 measurement system as reference is presented.

### 2.5 Cognitive tests

Five cognitive tests were conducted to identify differences in cognitive function between the three participant groups. As the current study is a cross-sectional comparison, it is impossible to show whether the continued practice of Wellness Darts in older adults affects cognitive function; however, it was preliminarily investigated to confirm its possibility. That is why the statistical tests were conducted between two groups: the expert older and the non-expert older groups.

#### 2.5.1 Simple Cognitive test

The Simple Cognitive (SC) test was used to verify that the participant’s cognitive function had not declined [27]. The SC test is a 50-point cognitive function test that can identify early cognitive decline that cannot be detected by the mini-mental state examination (MMSE), which is often used to examine dementia. A score of fewer than 20 points on the SC test is considered suspicious of cognitive decline. In this study, only participants with a score of 20 or higher were considered to have no cognitive decline and were used in subsequent analyses.

#### 2.5.2 Computational ability test

The participants’ computational abilities were measured by administering a test consisting of 30 subtraction problems between two digits, between three and two digits, and between three digits. Differences in mean scores on the computational ability test between the expert older group and the non-expert older group were *t*-tested (*p* < .05).

#### 2.5.3 STM

To evaluate the short-term memory of the participants, the Short-Term Memory (STM) Test was conducted on a computer [7]. After the instruction “Please memorize the numbers in order” was displayed on the monitor, nine disordered numbers were displayed on the screen one by one per second. The participants were then instructed to “write the numbers in order,” and they wrote the memorized numbers on a piece of paper during a 20-s time frame. The difference in mean STM test scores between the expert older group and the non-expert older group was *t*-tested (*p* < .05).

#### 2.5.4 CogEvo

CogEvo was used to test the cognitive abilities of the participants; CogEvo is a computer-assisted cognitive function test that assesses disorientation, attention, memory, planning, and spatial awareness [28]. CogEvo can reportedly capture mild changes in cognitive function and is a simple and convenient ICT tool to assess cognitive changes over time in middle-aged and older adults, as well as during the preclinical stages of dementia [28, 29]. CogEvo has 12 different tasks; in this experiment, we selected flashlight, visual search, number step, and just fit as tasks related to Wellness Darts from the disorientation, attention, memory, planning, and spatial awareness components of CogEvo, respectively. The difference in mean CogEvo scores between the expert older group and the non-expert older group was *t*-tested (*p* < .05).

#### 2.5.5 TMPT

The Trail making peg test (TMPT) was conducted to assess participants’ cognitive functioning, particularly attentional function and finger dexterity [30]. The TMPT is a peg-movement test that reflects finger dexterity combined with the trail-making test that measures visual attention and executive function [31]. In the TMPT, participants quickly insert 25 cylindrical wooden sticks, called pegs, into numbered holes in numerical order. The board containing the pegs and the board containing the holes were separated from each other. The difference in mean completion time was *t*-tested (*p* < .05).

### 2.6 Dart throwing performance test

To confirm that the dart-throwing skill of the expert older group was higher than that of the control groups, the dart-throwing performances of the three groups were tested. In this test, a target score of 65 points was set and participants were required to throw the darts 10 times and tally their scores to match the target score. The differences in total dart performance test scores between the expert and younger groups and between the expert and non-expert older groups were t-tested (*p* < .05).

### 2.7 Questionnaires

Participants were asked to respond to questions regarding their dominant hand, the number of days per week they practiced Wellness Darts, and the number of years they had been practicing. From these responses, the total number of days of experience for each participant was estimated.

### 2.8 fNIRS data processing

#### 2.8.1 Preprocessing

Previous studies have concluded that HbO in fNIRS is more strongly related to blood oxygenation level-dependent (BOLD) signals measured by functional magnetic resonance imaging (fMRI) than HbR [32, 33, 34]. The fMRI has better spatial resolution than fNIRS and has been used in many cognitive neuroscience studies. In this study, we used HbO to analyze brain activity. The fNIRS data were preprocessed using the NIRS toolbox [26] to remove physiological noise such as skin blood flow, heart rate, and pulse rate. Optical intensity data acquired from the fNIRS system were converted to optical density (OD) data and then to Hb concentration data based on the modified Beer-Lambert law [22]. A bandpass filter (0.008-0.09 Hz) was applied to remove noise such as low-frequency drift and heartbeat [34, 35].

#### 2.8.2 Activation analysis

The level of task-induced brain activation for each individual was estimated using the iteratively reweighted least-squares model (AR-IRLS) with short separation (SS) and acceleration, a general linear model (GLM)-based analysis method [24, 36]. In GLM, task-related brain activation was modeled by the hemodynamic response function (HRF). The measured brain activity, i.e., HbO signal, was regressed on the ideal task-related activity calculated from the canonical HRF and a design matrix indicating the type of experimental task, and the partial regression coefficients were estimated by IRLS, a least-squares method. The AR-IRLS allows brain activation analysis to consider the effects of arbitrary confounding factors on brain activity, such as the SS signal measured with a short-distance channel, blood pressure, respiration, and acceleration. In this study, we introduced the SS signal and acceleration data as physiological regressors of GLM to reduce the effects of skin blood flow and body motion on the HbO signal [24, 37].

#### 2.8.3 Hemispheric lateralization analysis

The level of hemispheric lateralization was calculated from the lateralization index (LI) shown in Equation 1.

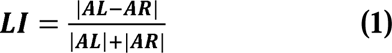

where AL is the sum of β values in the regions of interest (ROIs) in the left hemisphere and AR is the sum of those in the right hemisphere [38, 39]. We used absolute values for the LI values to allow for the fact that the dominant hemisphere differs depending on the dominant hand.

### 2.9 Statistical analysis

#### 2.9.1 Activation analysis

For individual-level analysis, the following two contrasts were created for the HbO data of the younger group, the expert older group, and the non-expert older group: “rest vs. calculation” and “rest vs. planning”. The β-values for each experimental condition and each channel were calculated. The statistical significance of the task-related brain activity was then verified by testing the null hypothesis that the estimated β coefficient was not significantly different from zero (*t*-test, *p* < .05). Note that multiple comparisons were corrected using the Benjamini-Hochberg method [37]. In addition, to test the hypothesis that there is no difference in the level of IPL activation between older adults who continually practiced Wellness Darts and younger adults who were not experienced in Wellness Darts, β values of IPL were compared between the three groups (*t*-test, *p* < .016, Bonferroni corrected).

Next, for the group-level analysis, we constructed the most consistent contrast images of group-level brain activity using simultaneously the β values obtained from the individual-level analysis of all participants, based on [25]. To estimate significant group-level activity, we used participants as random effects and conditions (planning, calculation, and rest blocks) as fixed effects. Finally, all participants were considered as a sample drawn from the population and a one-sample t-test was used to test if the population mean was greater than zero with respect to the *t*-value. The significance level was set at *p* < .05 (false discovery rate corrected at peak level), and the brain regions activated during the calculation and planning blocks were estimated.

#### 2.9.2 Hemispheric lateralization analysis

The LI values were used to compare the degree of hemispheric lateralization among the three groups of participants (*t*-test, *p* < .016, Bonferroni corrected). In addition, we tested whether there was a positive correlation between the number of days of Wellness Dart experience and LI values in the expert older group (Spearmans’ rank correlation test, *p* < .05).

## 3 Results

### 3.1 Descriptive statistics on three groups

The descriptive statistics on three groups, including age, days of practicing wellness darts, cognitive test scores, and scores of dart-throwing performance tests, are summarized in Table 1. The following subsections describe the details of the results including statistical tests.

**Table 1.** Descriptive statistics on the three groups.

| Characteristics |  | Expert older group | Nonexpert older group (older control) | Younger group (nonexpert younger control) |
| --- | --- | --- | --- | --- |
| age (years) |  | 75.1±4.3 | 74.1±4.5 | 22.5±0.8 |
| Number of days of Wellness Darts experience (days) |  | 170.7±145.6 | 0.0 | 0.0 |
| Cognitive tests | SC test | 41.1±7.8 | 44.5±5.8 | 48.8±2.5 |
|  | Computational ability test | 20.5±3.8 | 17.1±5.5 | 24.0±3.8 |
|  | STM test | 57.1±9.5 | 49.4±9.9 | 74.6±13.6 |
|  | TMPT | 58.1±8.5 | 58.9±9.2 | 41.1±4.5 |
|  | CogEvo: disorientation | 361.8±52.7 | 278.2±74.5 | 341.5±68.8 |
|  | CogEvo: visual search | 177.9±39.0 | 161.3±33.8 | 282.0±41.3 |
|  | CogEvo: flashlight | 364.3±146.9 | 308.1±106.4 | 604.3±169.2 |
|  | CogEvo: number step | 197.4±40.8 | 173.5±45.9 | 280.9±42.8 |
|  | CogEvo: just fit | 332.2±104.9 | 342.4±98.7 | 442.1 ± 78.2 |
| Dart-throwing performance |  | 49.9±5.5 | 31.9±13.7 | 39.3±13.4 |

### 3.2 fNIRS data

#### 3.2.1 Hemispheric lateralization analysis

We hypothesized that the LI values of the younger group and the expert older group would be larger than those of the non-expert older group. LI values were calculated from the respective β values of the two task blocks (planning and calculation) for the three participant groups. The LI values in the planning and calculation blocks of the younger group were 0.45 ± 0.34 and 0.49 ± 0.35, respectively. The LI values in the planning and calculation blocks of the expert older group were 0.53±0.38 and 0.61±0.35, respectively. Finally, for the non-expert older group, they were 0.23±0.18 and 0.50±0.38 in the planning and calculation blocks, respectively. The box plots of the LI values in the planning and calculation blocks for the three groups of participants are shown in Figure 3 (d) and Figure 4 (d), respectively. Furthermore, the β maps of the participants who showed the closest values to the median in the LI values for each group of participants are shown in Figure 3 (a)-(c) and Figure 4 (a)-(c).

**Figure 3.**
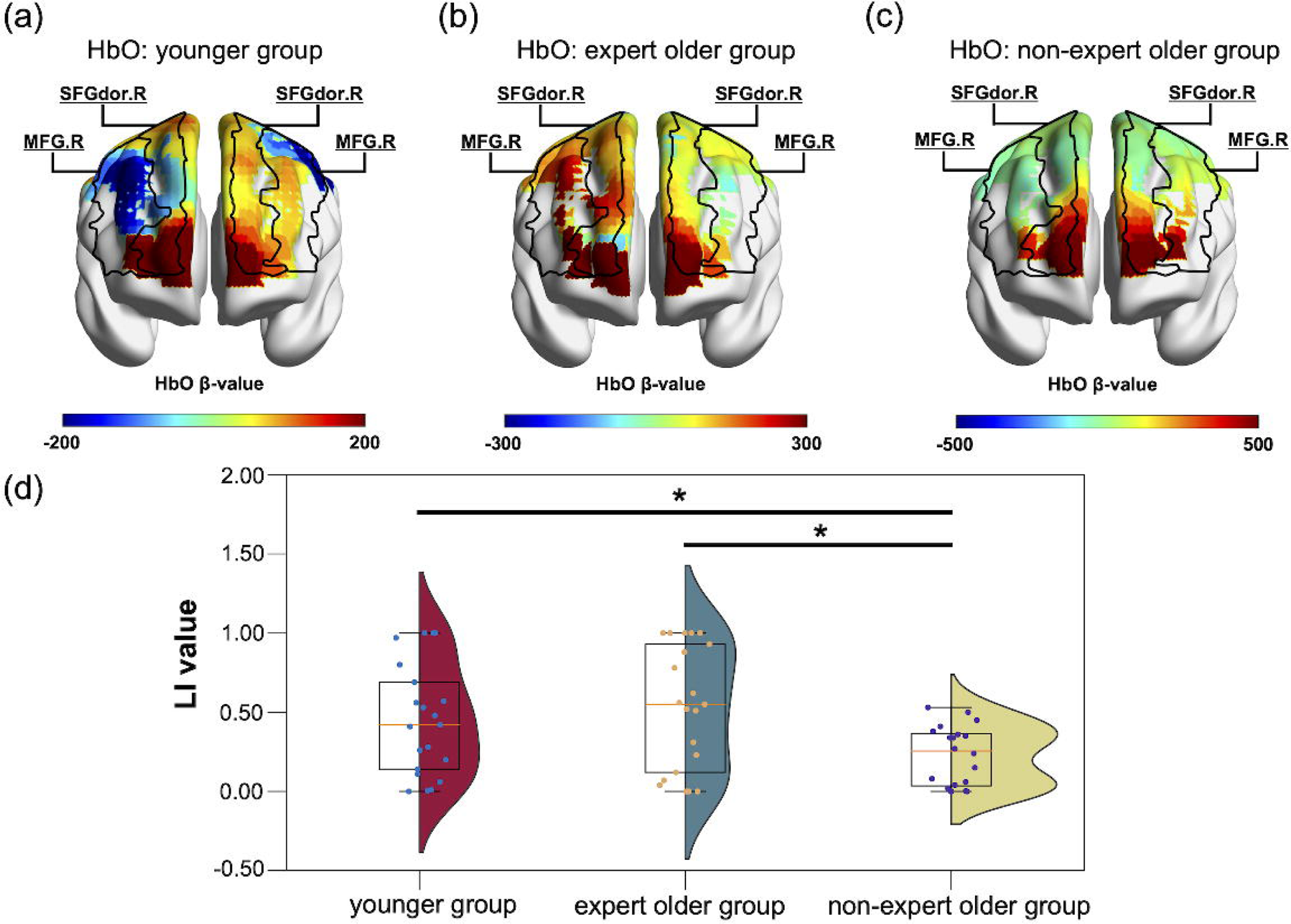
βmaps and LI values in the planning block. (a)-(c) βmaps of participants with LI values close to the median in younger, expert older, and non-expert older groups are shown, respectively. (d) Distribution of LI values for the three participant groups. *p < .016. βmaps were visualized using the xjView toolbox and BrainNet viewer.

**Figure 4.**
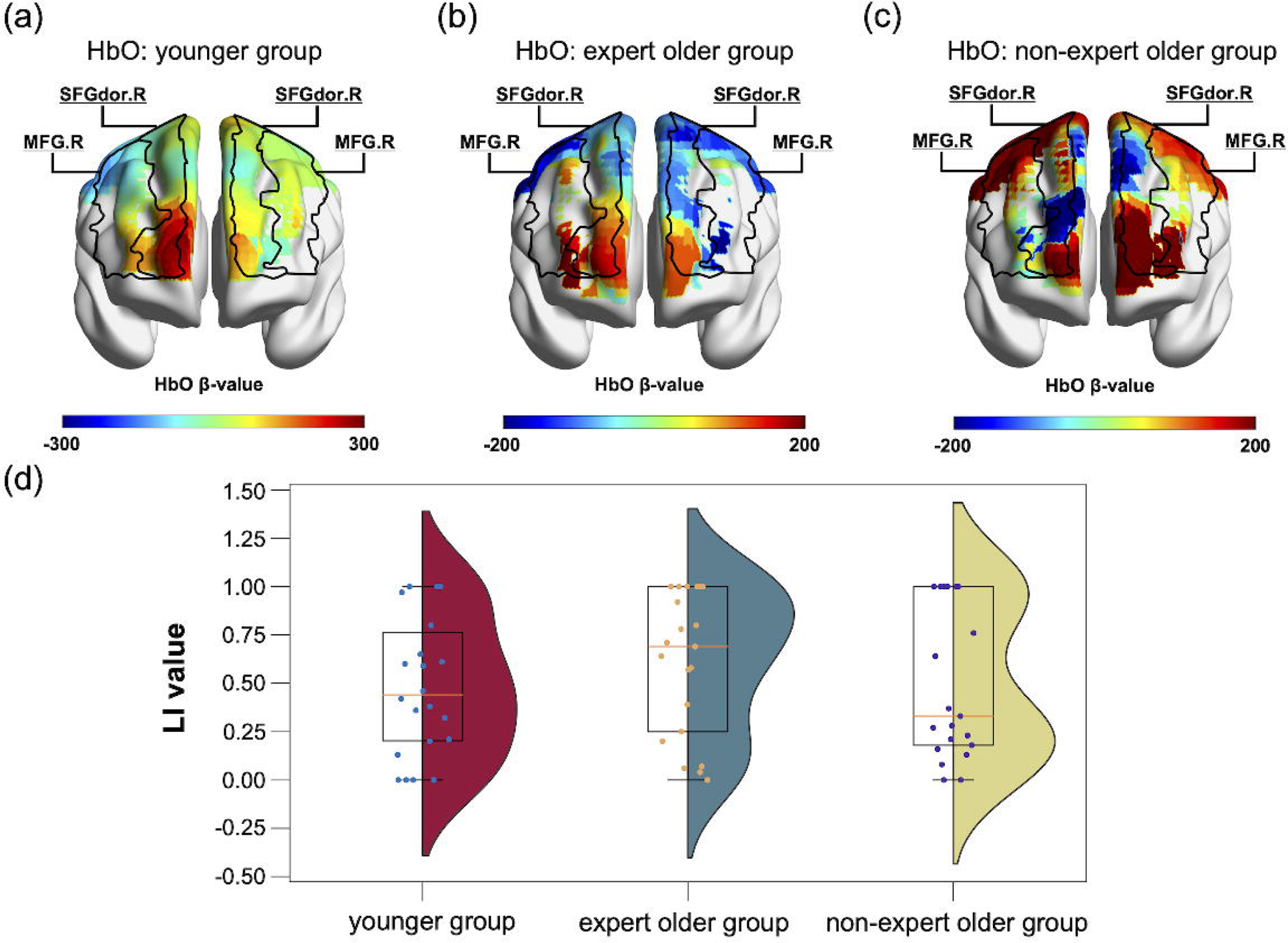
βmaps and LI values in the calculation block. (a)-(c) βmaps of participants with LI values close to the median in the younger, expert older, and non-expert older groups are shown, respectively. (d) Distribution of LI values for the three participant groups. *p < .016. βmaps were visualized using the xjView toolbox and BrainNet viewer.

There were no significant differences between the LI values of the younger group and the expert older group in the planning block (*t* = 0.6730, *p* = 0.504, *d* = 0.213). On the other hand, in the planning block, the LI values of the expert older group were significantly greater than those of the non-expert older group (*t* = 3.188, *p* = 0.003, *d* = 1.005). Furthermore, the LI values of the younger group were significantly greater than those of the non-expert older group in the planning block (*t* = 2.58, *p* = 0.014, *d* = 0.815).

In the calculation block, there were no differences in LI values between the younger group and the expert older group (*t* = 1.009, *p* = 0.319, *d* = 0.319). Similarly, there were no differences in LI between the expert older group and the non-expert older group (*t* = 0.828, *p* = 0.413, *d* = 0.262). There were also no differences in LI values between the younger group and the non-expert older group (*t* = 0.144, *p* = 0.886, *d* = 0.046).

We also hypothesized that the number of days of Wellness Dart experience in the expert older group would be positively correlated with LI values. The results of an Spearman’s rank correlation test between the number of days of Wellness Dart experience and LI values showed a negative significant correlation for the planning block (ρ = −0.439, *p* = 0.046, Figure 5 (a)) but no significant correlation for the calculation block (ρ = −0.037, *p* = 0.874, Figure 5 (b)).

**Figure 5.**
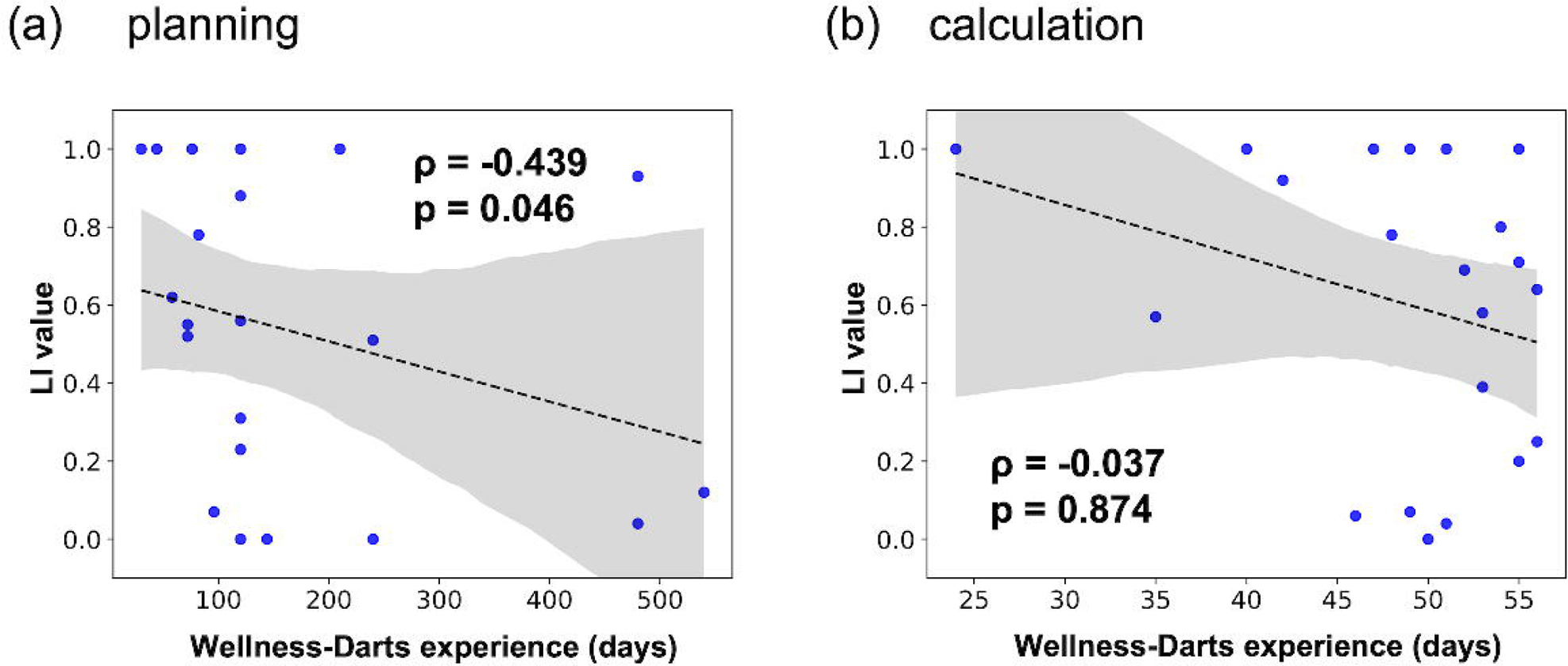
Correlation between the number of days of Wellness Dart experience and LI values in the expert older group. (a) planning block. (b) calculation block.

To investigate the Wellness Dart experience of the expert older group in terms of skill, the mean darts performance test score (target score was 65 points; the closer the total score of 10 dart throws was to 65 points, the better the skill) was compared between the three participant groups. The mean scores for the younger group, the expert older group, and the non-expert older group were 39.3±13.4, 49.9±5.5, and 31.9±13.7, respectively. There were no participants who exceeded the 65 points. As shown in Figure 6, the expert older group showed better dart performance compared to the younger group (*t* = 3.247, *p* = 0.003, *d* = 1.022). Similarly, the expert older group showed better performance than the non-expert older group (*t* = 5.400, *p* = 0.00001, *d* = 1.700).

**Figure 6.**
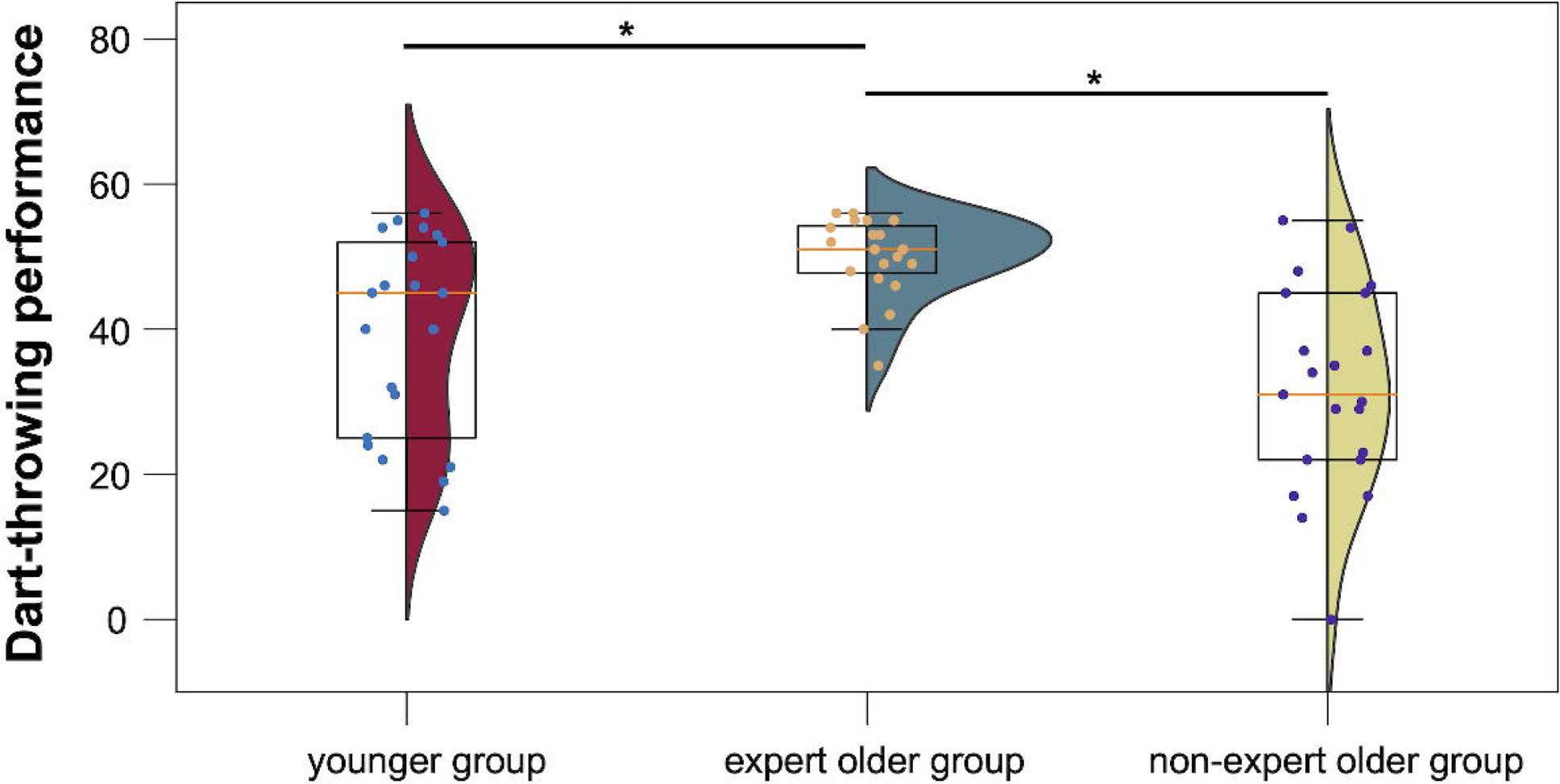
Dart performance level in the younger, expert older, and non-expert older group. *p<.05.

Furthermore, inspired by the results of no significant ‘positive’ correlation between the number of days of Wellness Dart experience and LI values, we assumed that dart proficiency could be assessed not only by the number of days of experience but also by dart-throwing performance. Therefore, a positive correlation between the total dart performance test score and the degree of hemispheric lateralization of brain activity in the expert older group was also tested (Spearman’s rank correlation test, *p* < .05). It should be noted that this is an exploratory analysis based on the result shown in Figure 5. However, no significant correlation between the total dart performance test score and the LI value was confirmed (planning block: ρ = 0.324, *p* = 0.152, Figure 7 (a); calculation block: ρ = −0.225, *p* = 0.327, Figure 7 (b)).

**Figure 7.**
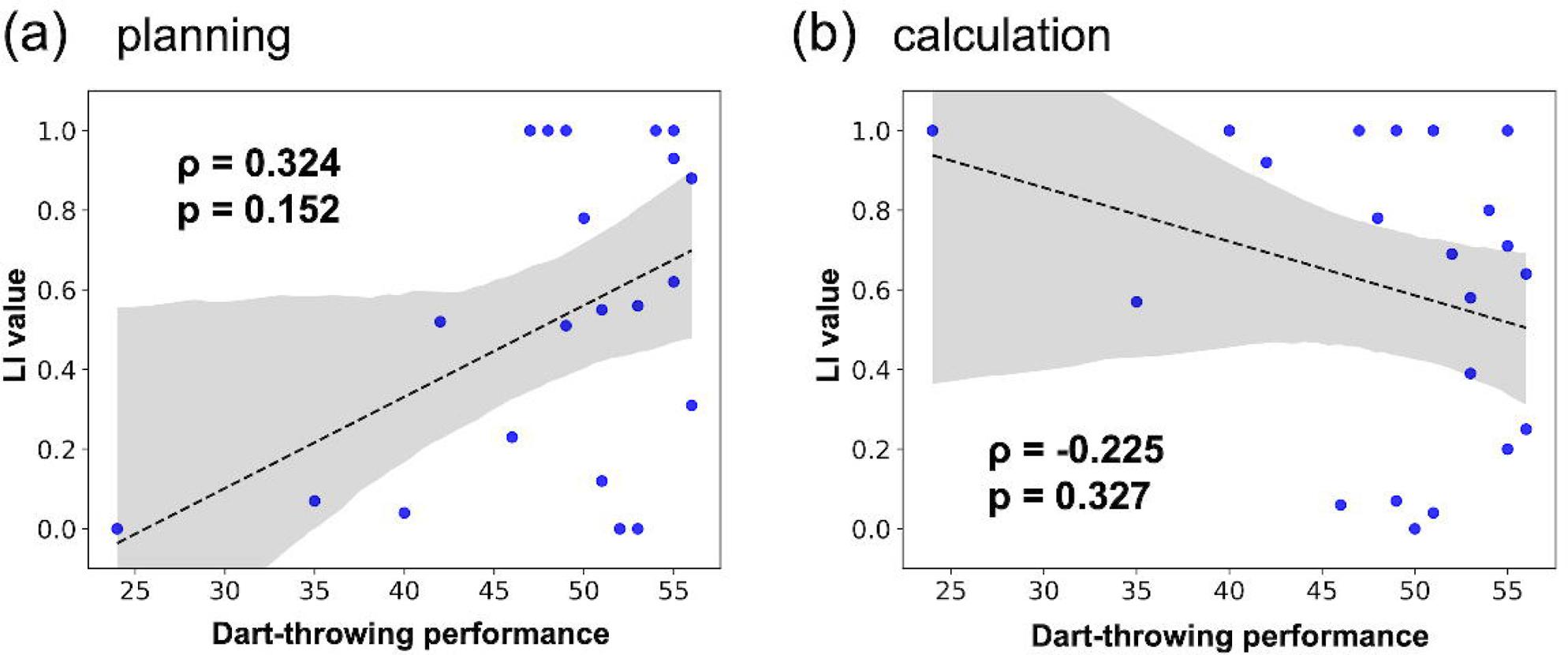
Correlation between darts performance level and LI value in the expert older group. (a) planning block. (b) calculation block.

#### 3.2.2 Activation analysis

The activation analysis shown in Figure 8 indicates that in the planning block, there was no difference between the left and right IPL (IPL.L: *t* = 0.738, *p* = 0.466, *d* = 0.234. IPL.R: *t* = 0.134, *p* = 0.894, *d* = 0.043). Similarly, there was no difference between the expert older group and the non-expert older group (IPL.L: *t* = 0.042, *p* = 0.966, *d* = 0.014; IPL.R: *t* = 0.821, *p* = 0.419, *d* = 0.257). Furthermore, there was no difference between the younger group and the non-expert older group (IPL.L: *t* = 0.711, *p* = 0.481, *d* = 0.226. IPL.R: *t* = 0.967, *p* = 0.343, *d* = 0.312).

**Figure 8.**
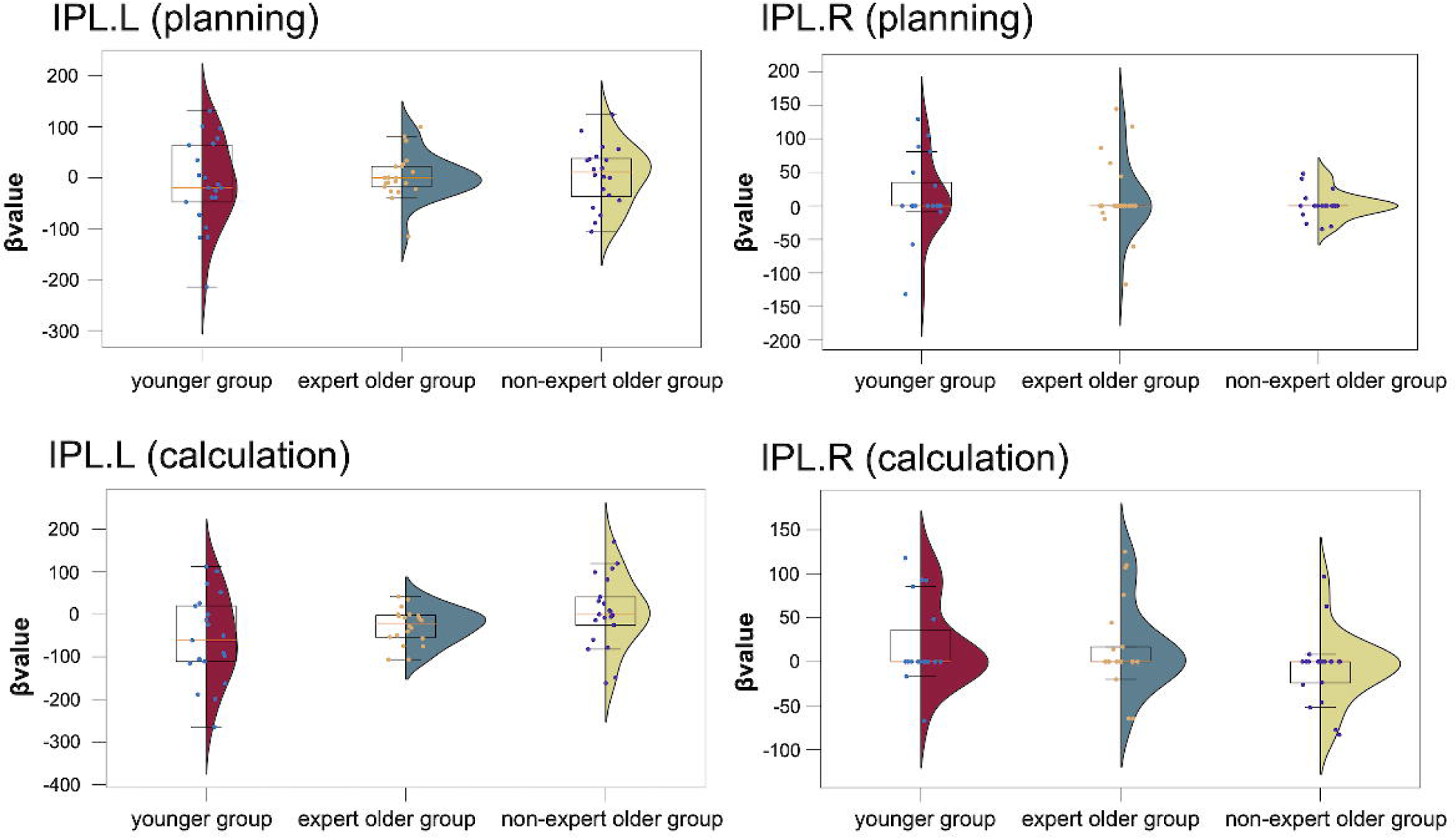
β values of IPL in the younger, expert older, and non-expert older group.

In the calculation block, there was no difference between the left and right IPL (IPL.L and IPL.R) of the younger group and the expert older group (IPL.L: *t* = 1.211, *p* = 0.236, *d* = 0.381; IPL.R: *t* = 0.206, *p* = 0.838, *d* = 0.068). Similarly, there was no difference between the expert older group and the non-expert older group (IPL.L: *t* = 1.625, *p* = 0.115, *d* = 0.513; IPL.R: *t* = 1.642, *p* = 0.109, *d* = 0.523). There was no difference between the younger group and the non-expert older group (IPL.L: *t* = 2.176, *p* = 0.036, *d* = 0.688. IPL.R: *t* = 1.844, *p* = 0.074, *d* = 0.621).

Tables 2 to 7 summarize the results of individual-level analysis and show the regions of interest where the *p*-values are less than 0.05. Tables 8 to 10 summarize the results of group-level analysis and show only the ROIs. There was significant activation in the ROIs in the individual-level analysis in the expert older group but no significant activation in the group-level analysis.

**Table 2.**
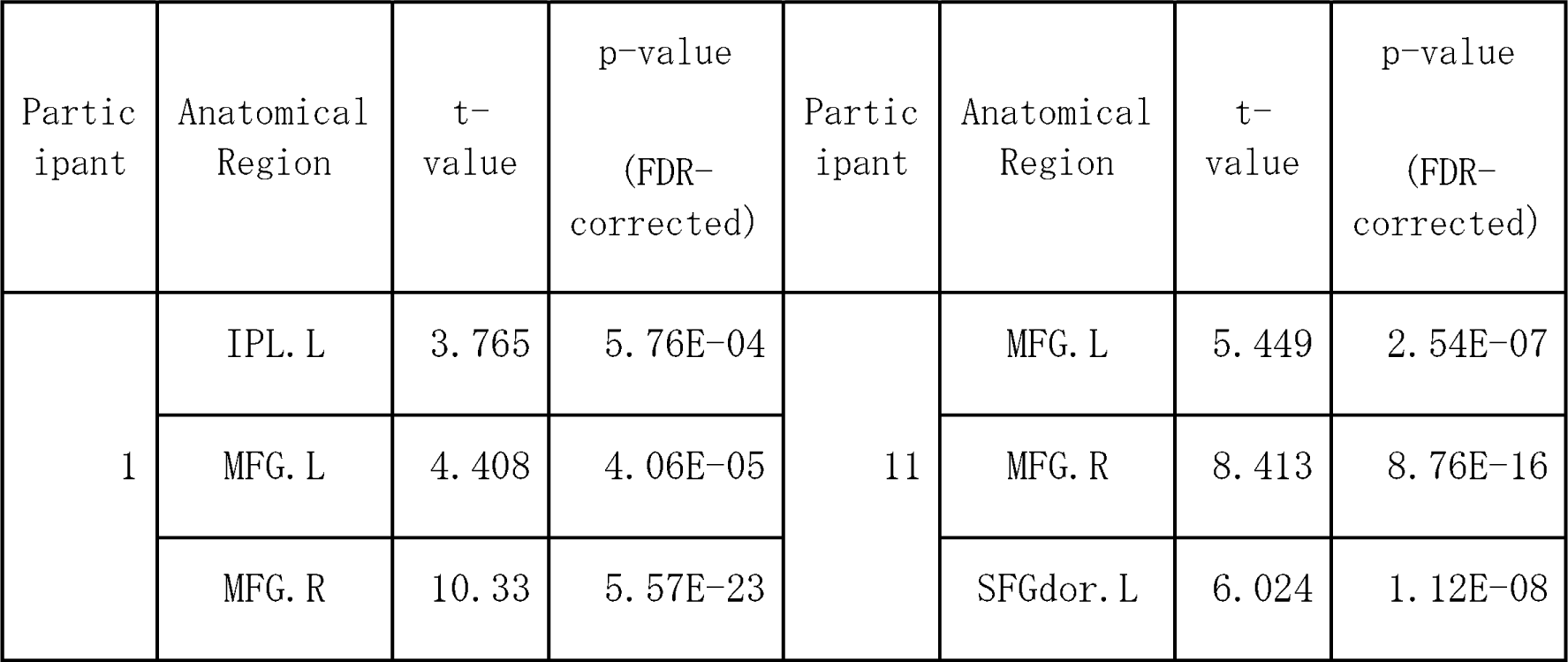
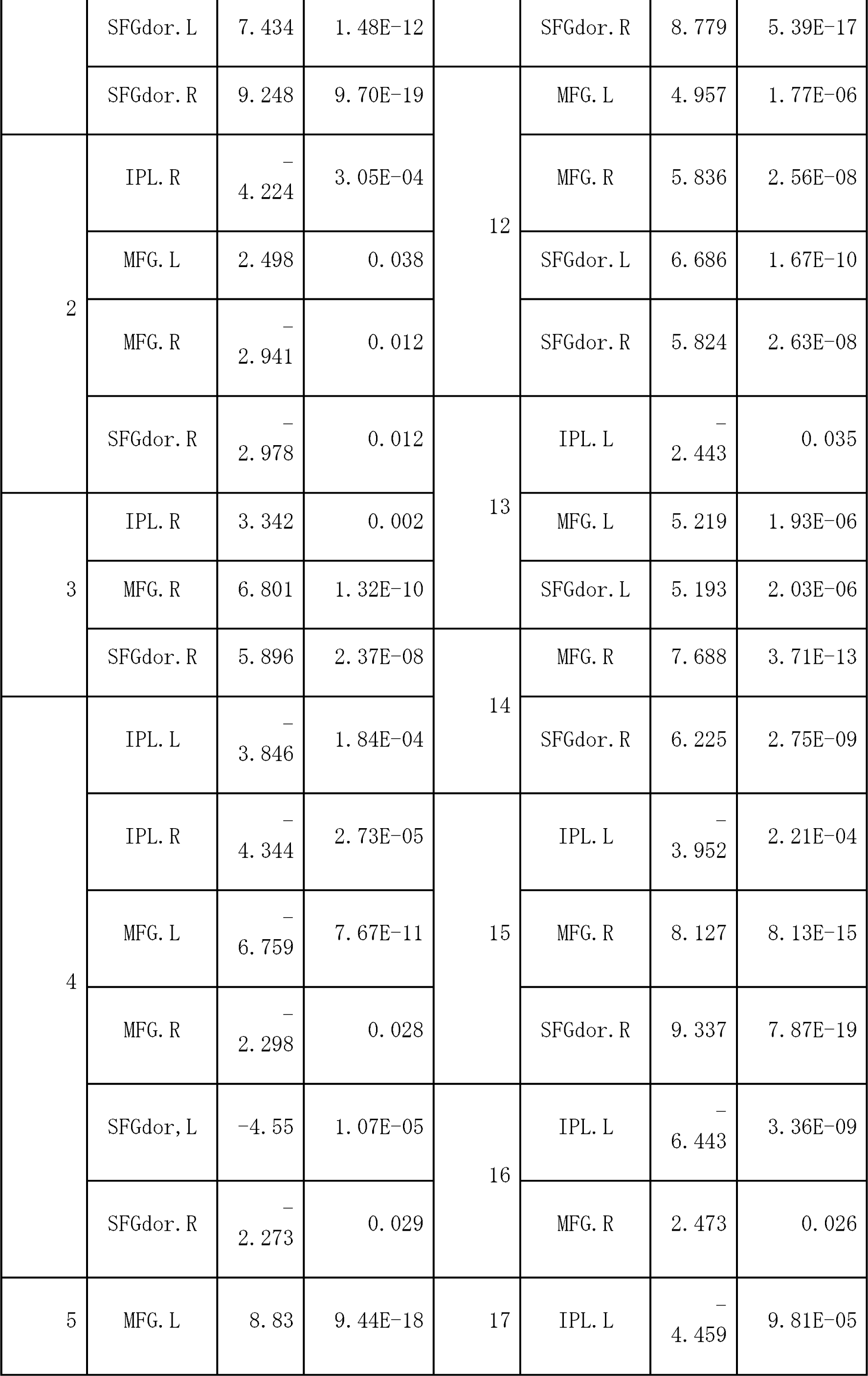
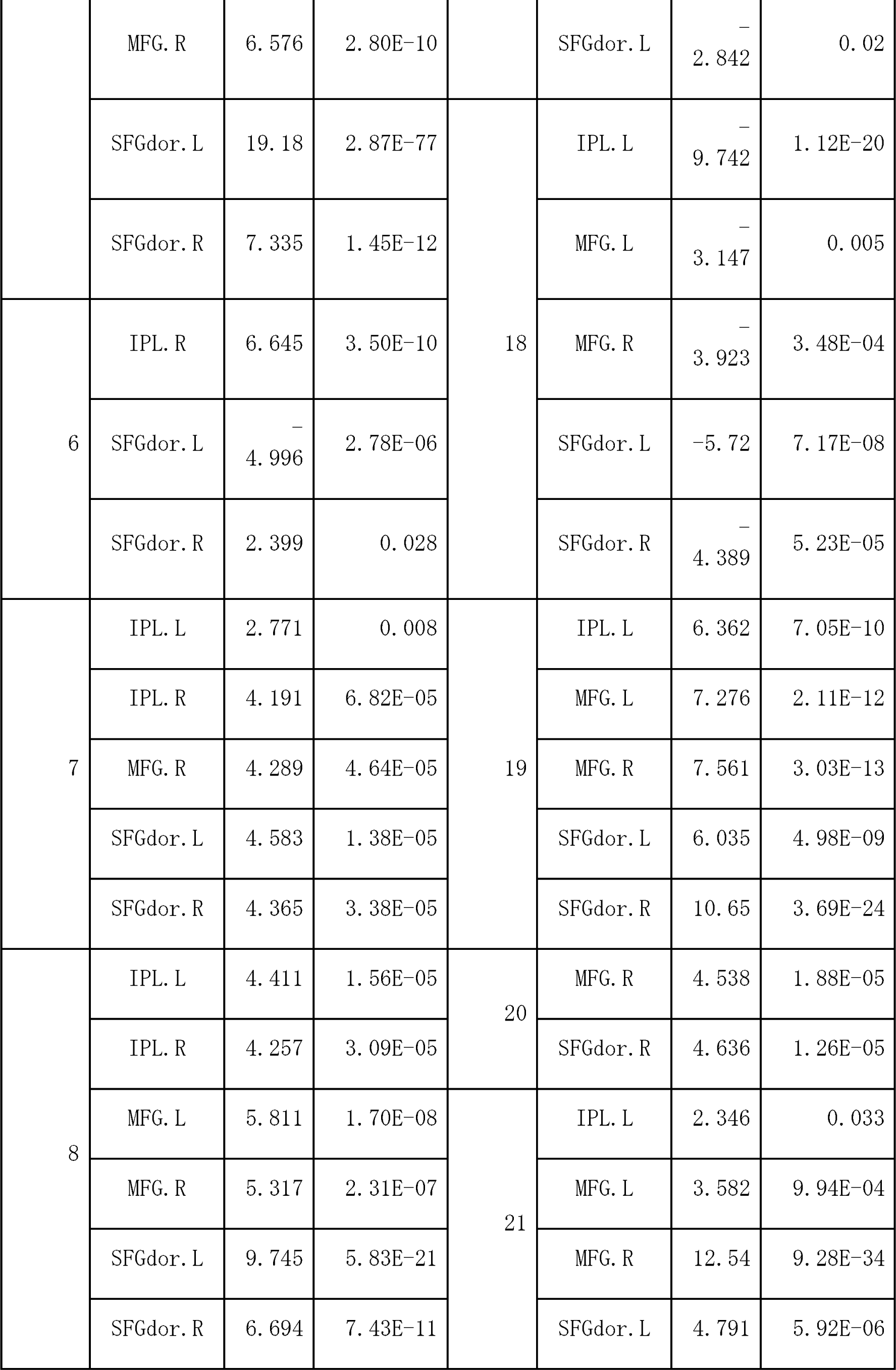
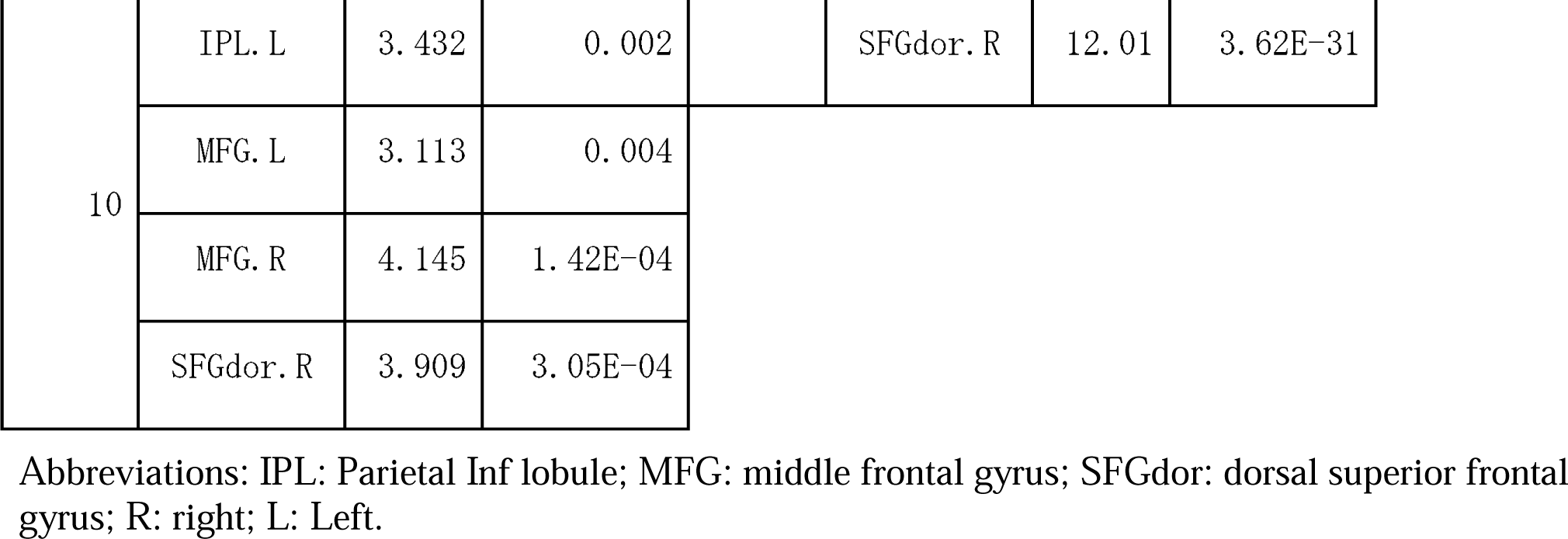
Brain regions that showed task-related activation for each individual in the planning block (younger group). Only regions of interest that showed p < 0.05 (FDR corrected at peak level) are shown.

**Table 3.**
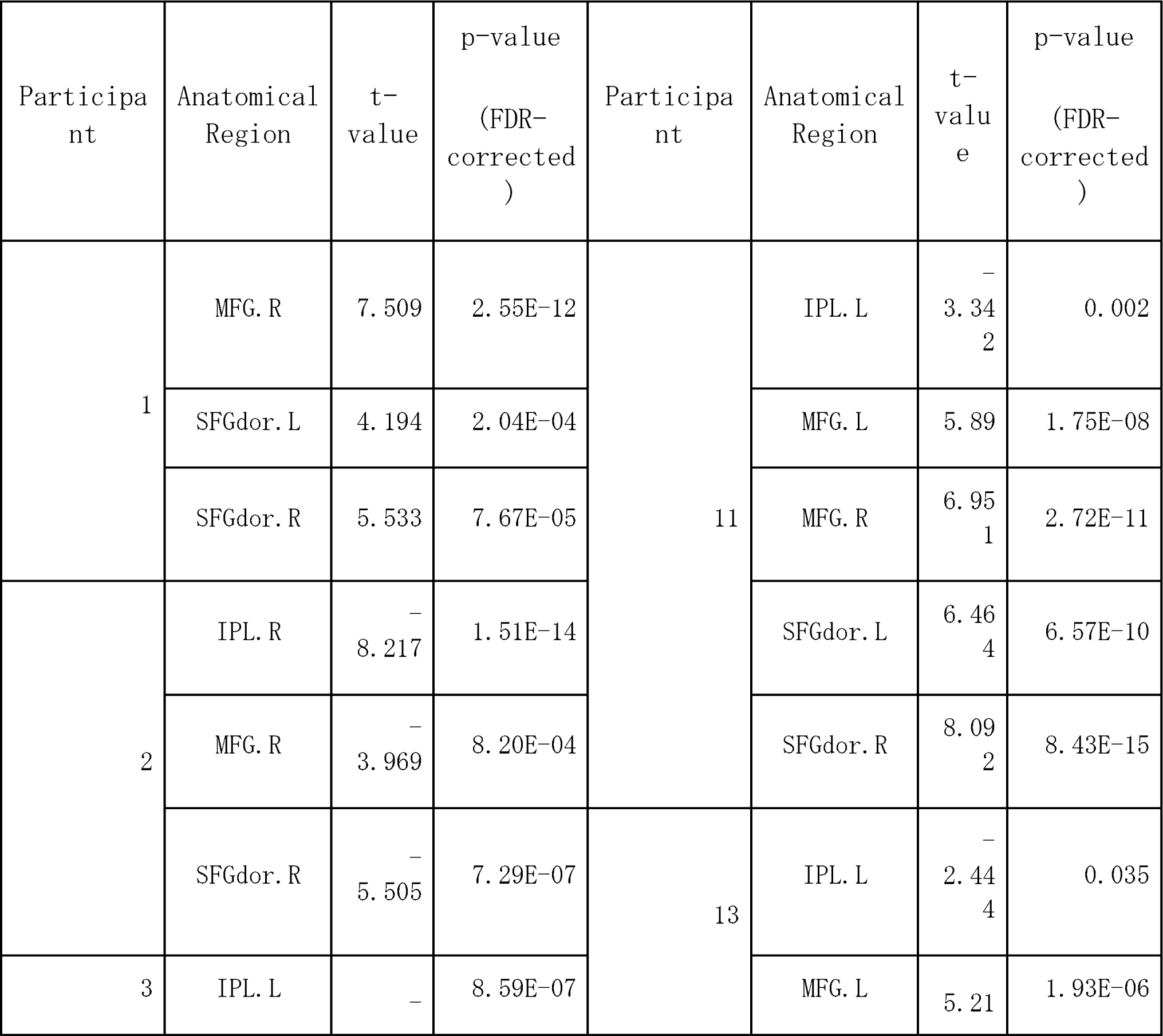
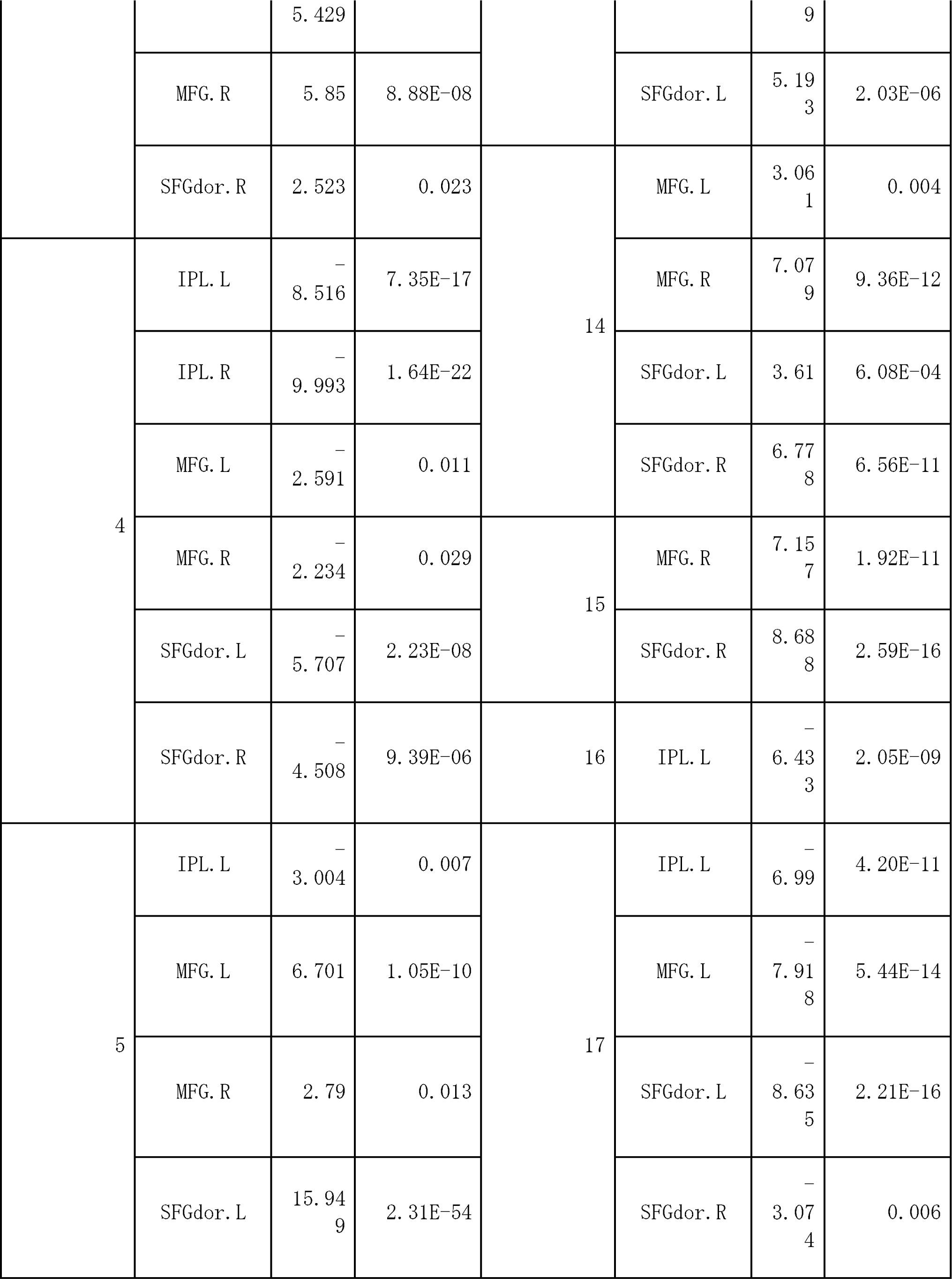
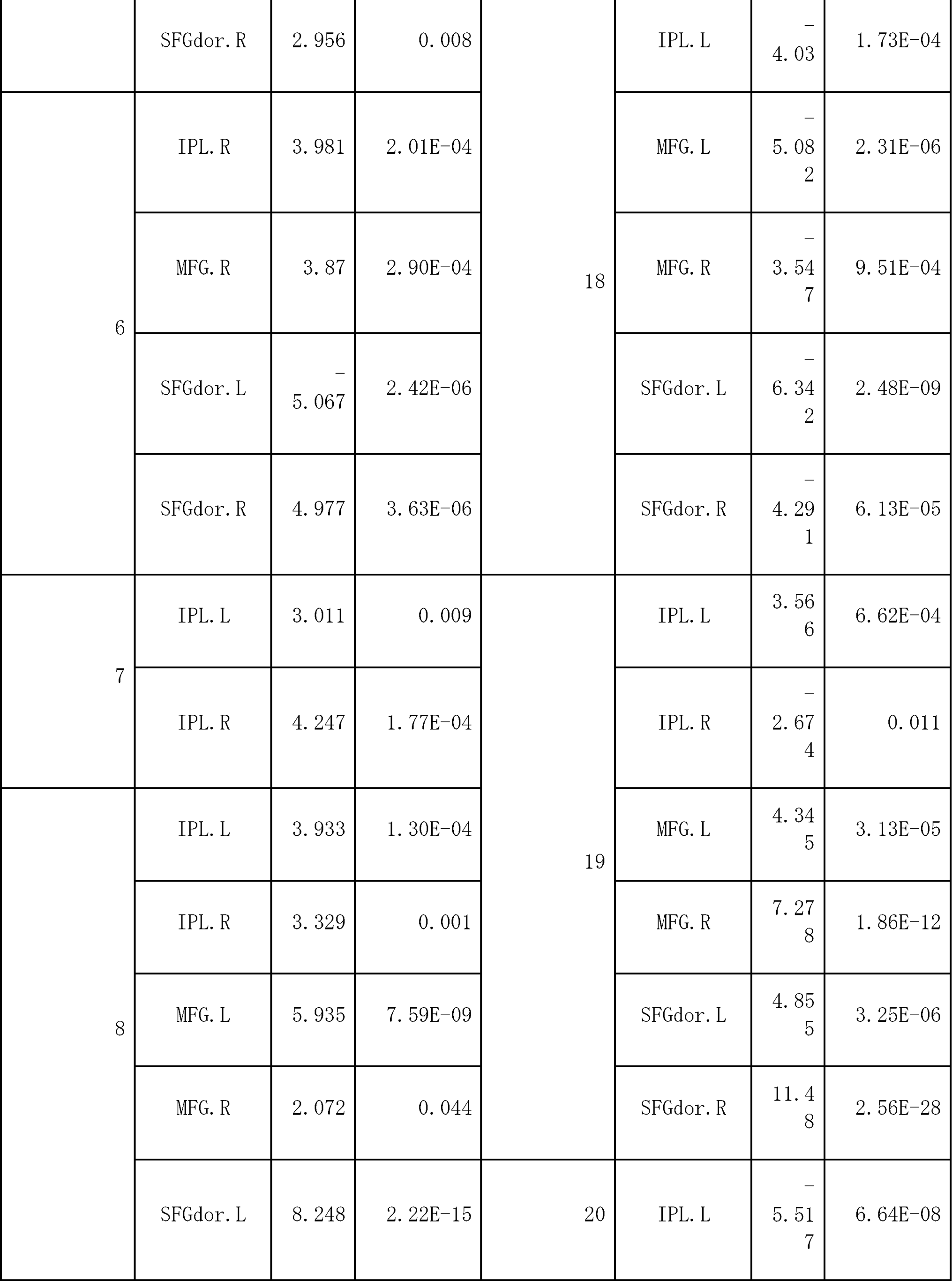
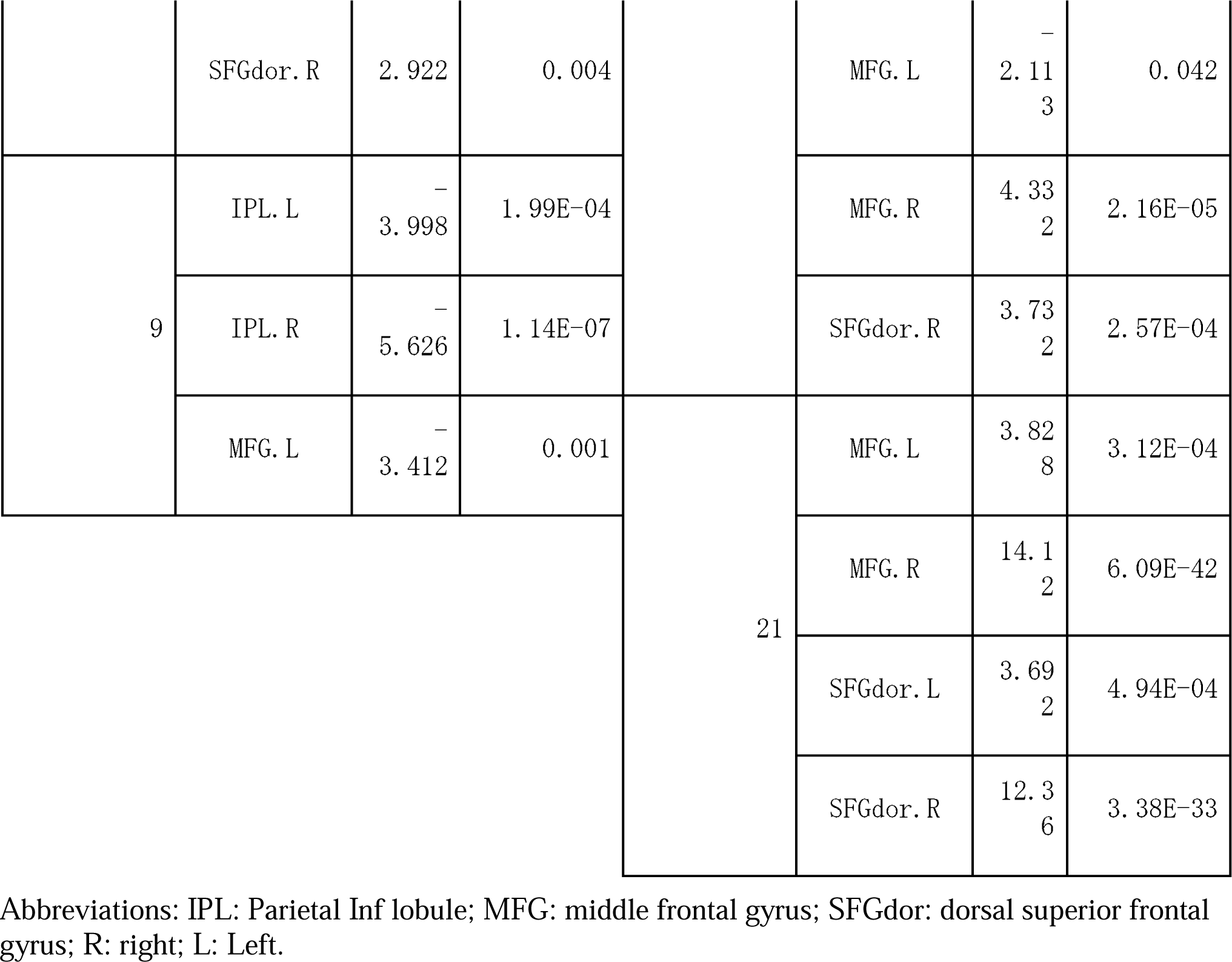
Brain regions that showed task-related activation for each individual in the calculation block (younger group). Only regions of interest that showed p < 0.05 (FDR corrected at peak level) are shown.

**Table 4.**
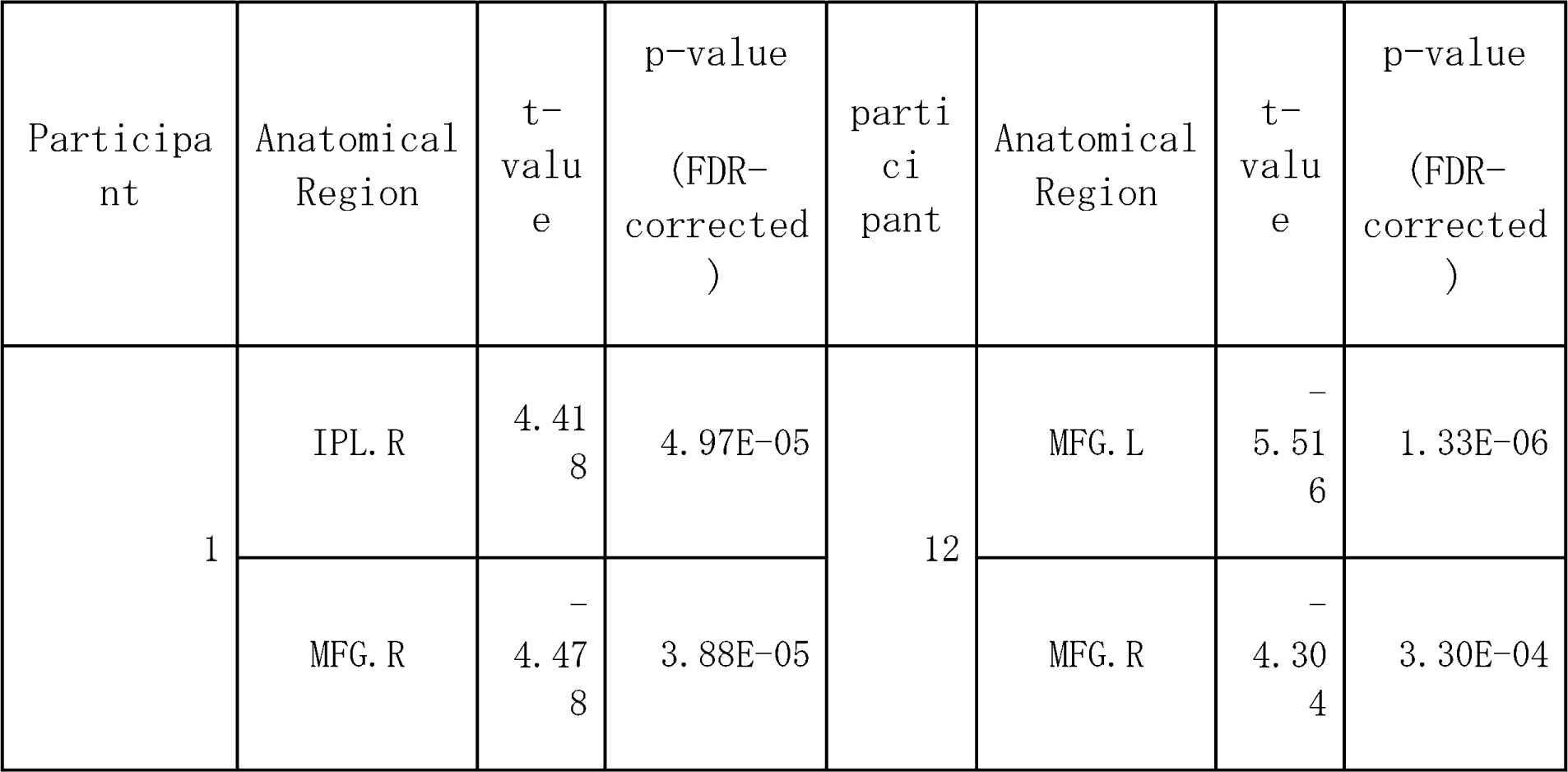
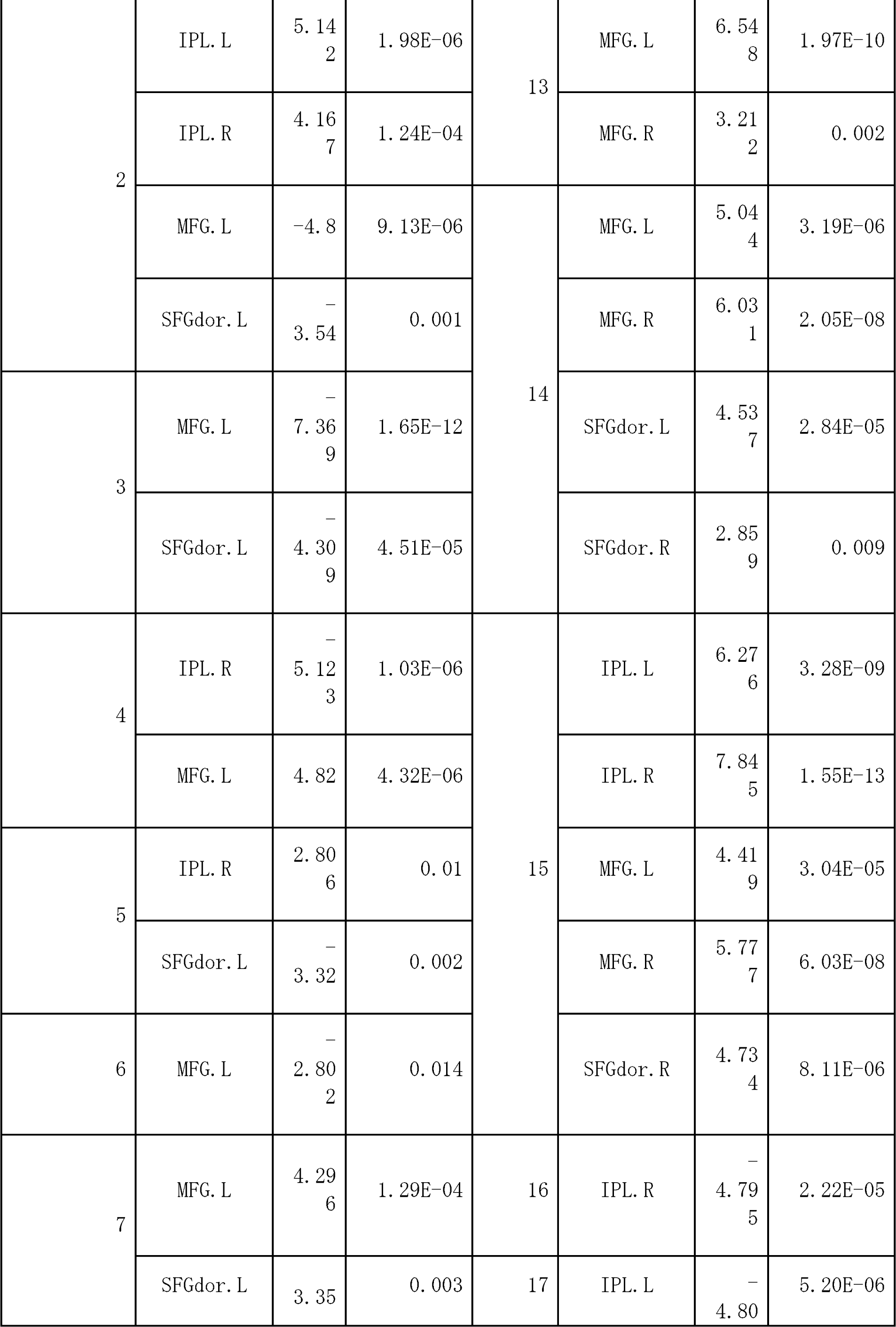
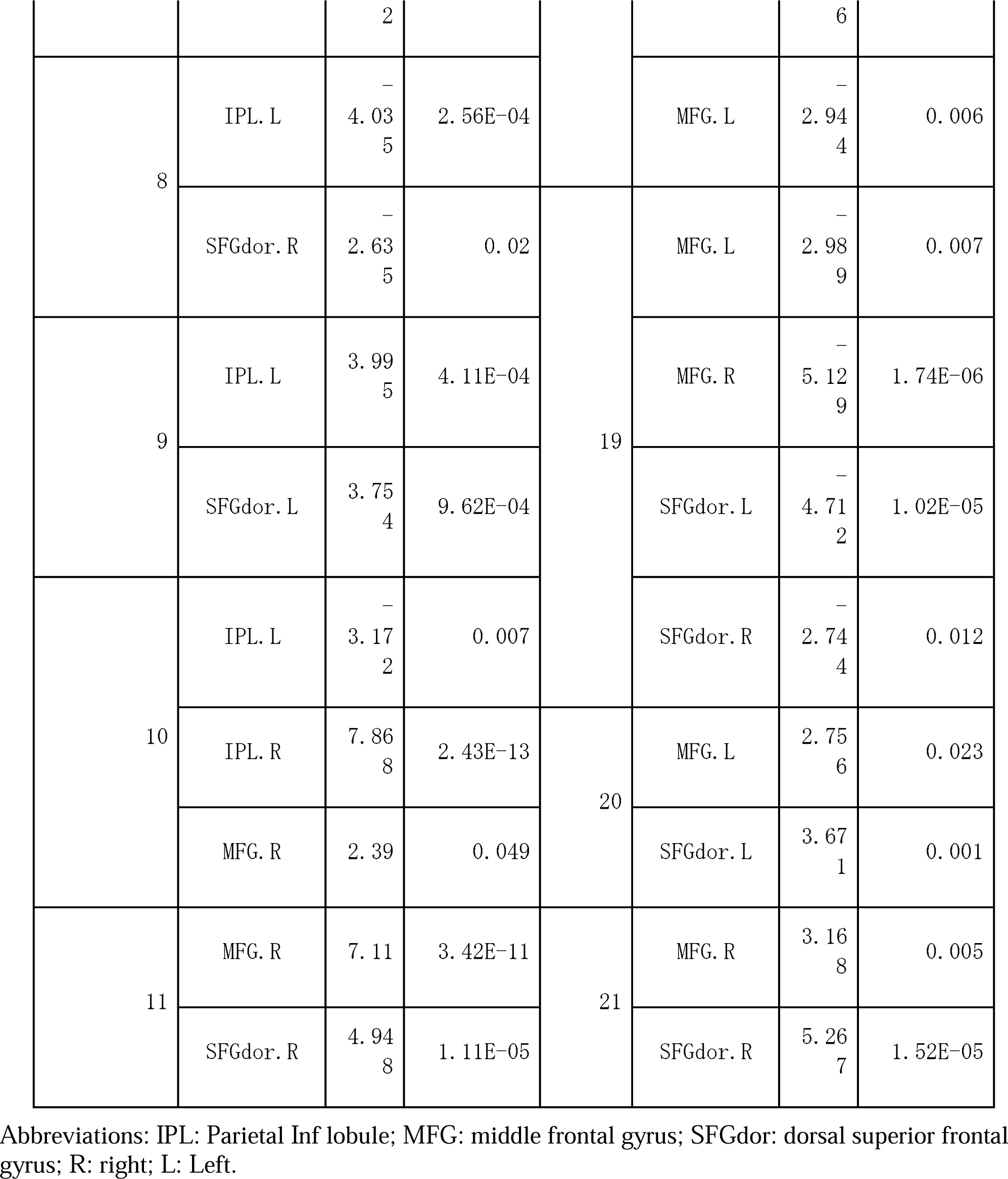
Brain regions that showed task-related activation in the planning block for each individual (expert older group). Only regions of interest with p < 0.05 (FDR corrected at peak level) are shown.

**Table 5.**
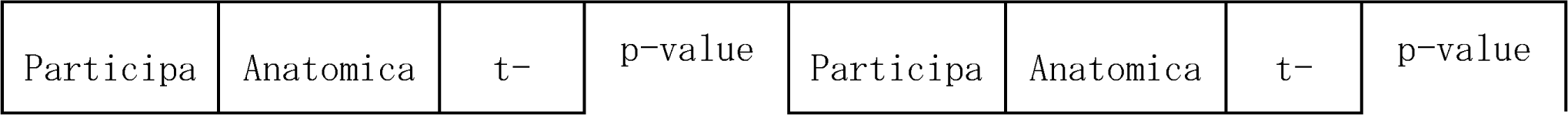
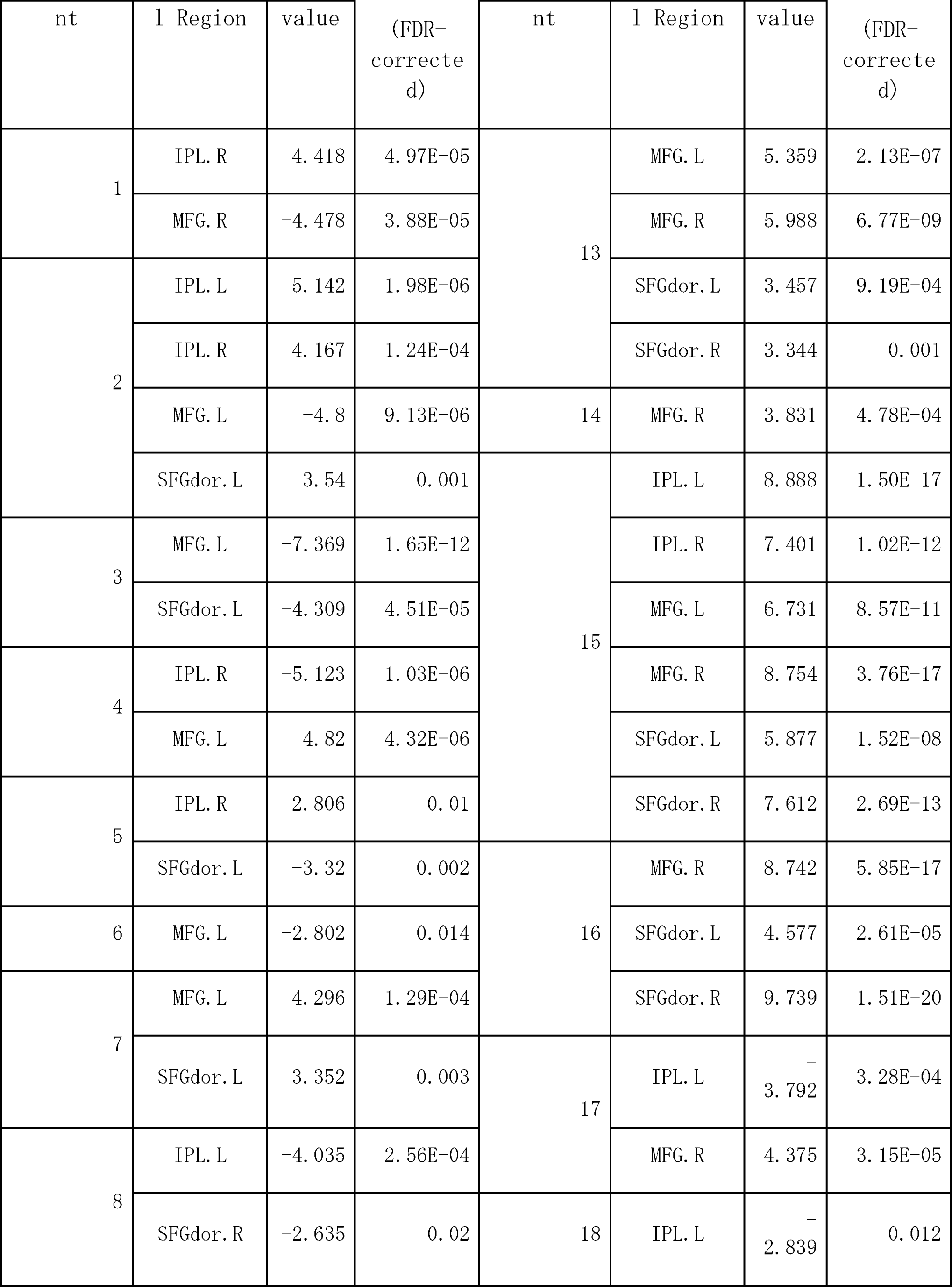
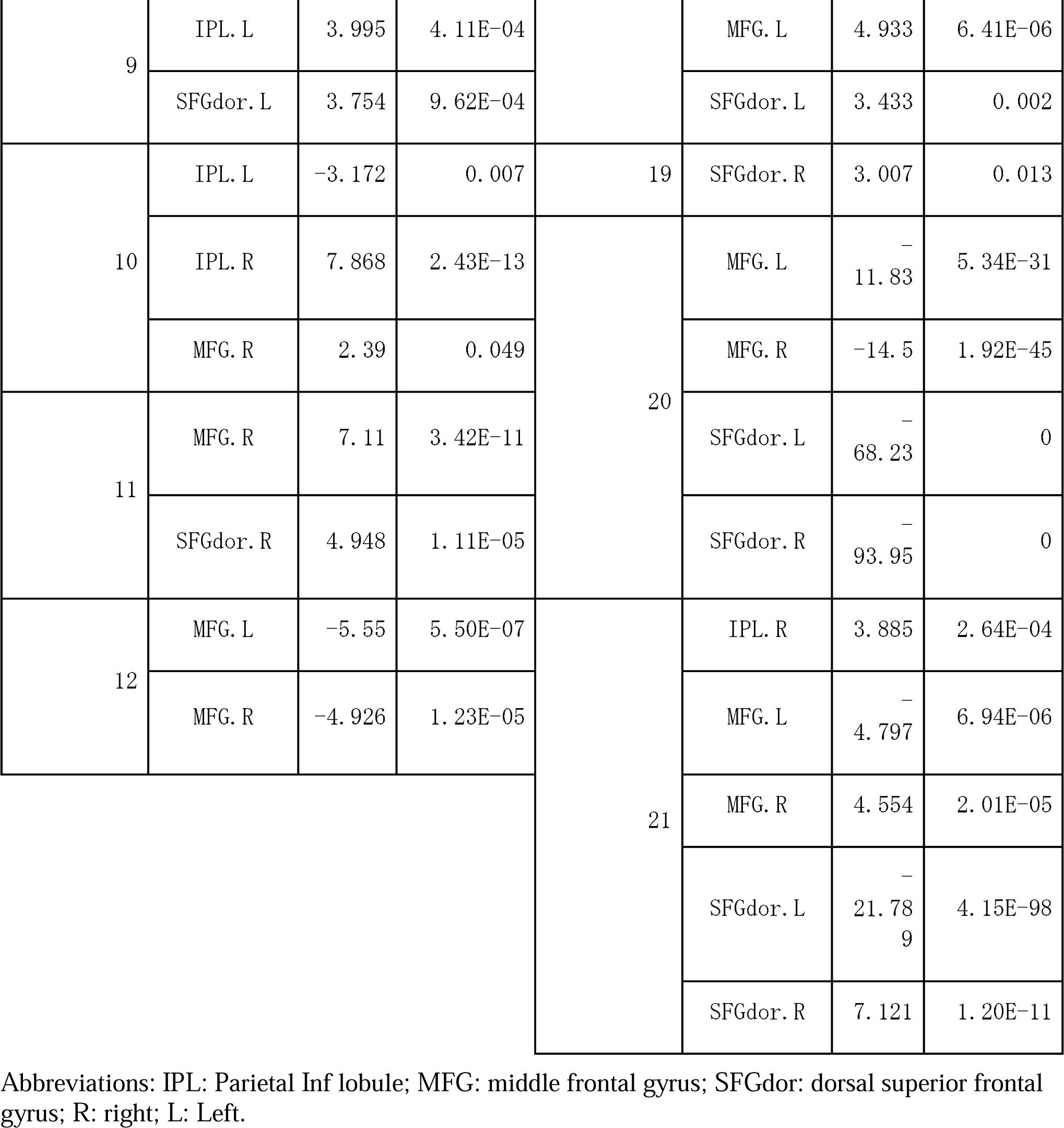
Brain regions that showed task-related activation for each individual in the calculation block (expert older group). Only regions of interest with p < 0.05 (FDR corrected at peak level) are shown.

|  |  |  |  |  |  |  |  |
| --- | --- | --- | --- | --- | --- | --- | --- |
| Participa | Anatomica | t- | p-value | Participa | Anatomica | t- | p-value |

Hemispheric lateralization in dart players
| nt | l Region | value | (FDR-corrected) | nt | l Region | value | (FDR-corrected) |
| --- | --- | --- | --- | --- | --- | --- | --- |
| 1 | IPL. R | 4.418 | 4.97E-05 | 13 | MFG. L | 5.359 | 2.13E-07 |
|  | MFG. R | -4.478 | 3.88E-05 |  | MFG. R | 5.988 | 6.77E-09 |
| 2 | IPL. L | 5.142 | 1.98E-06 |  | SFGdor. L | 3.457 | 9.19E-04 |
|  | IPL. R | 4.167 | 1.24E-04 |  | SFGdor. R | 3.344 | 0.001 |
|  | MFG. L | -4.8 | 9.13E-06 | 14 | MFG. R | 3.831 | 4.78E-04 |
|  | SFGdor. L | -3.54 | 0.001 | 15 | IPL. L | 8.888 | 1.50E-17 |
| 3 | MFG. L | -7.369 | 1.65E-12 |  | IPL. R | 7.401 | 1.02E-12 |
|  | SFGdor. L | -4.309 | 4.51E-05 |  | MFG. L | 6.731 | 8.57E-11 |
| 4 | IPL. R | -5.123 | 1.03E-06 |  | MFG. R | 8.754 | 3.76E-17 |
|  | MFG. L | 4.82 | 4.32E-06 |  | SFGdor. L | 5.877 | 1.52E-08 |
| 5 | IPL. R | 2.806 | 0.01 |  | SFGdor. R | 7.612 | 2.69E-13 |
|  | SFGdor. L | -3.32 | 0.002 | 16 | MFG. R | 8.742 | 5.85E-17 |
| 6 | MFG. L | -2.802 | 0.014 |  | SFGdor. L | 4.577 | 2.61E-05 |
| 7 | MFG. L | 4.296 | 1.29E-04 |  | SFGdor. R | 9.739 | 1.51E-20 |
|  | SFGdor. L | 3.352 | 0.003 | 17 | IPL. L | -3.792 | 3.28E-04 |
| 8 | IPL. L | -4.035 | 2.56E-04 |  | MFG. R | 4.375 | 3.15E-05 |
|  | SFGdor. R | -2.635 | 0.02 | 18 | IPL. L | -2.839 | 0.012 |
| 9 | IPL. L | 3. 995 | 4. 11E-04 |  | MFG. L | 4. 933 | 6. 41E-06 |
|  | SFGdor. L | 3. 754 | 9. 62E-04 |  | SFGdor. L | 3. 433 | 0. 002 |
| 10 | IPL. L | -3. 172 | 0. 007 | 19 | SFGdor. R | 3. 007 | 0. 013 |
|  | IPL. R | 7. 868 | 2. 43E-13 | 20 | MFG. L | -<br>11. 83 | 5. 34E-31 |
|  | MFG. R | 2. 39 | 0. 049 |  | MFG. R | -14. 5 | 1. 92E-45 |
| 11 | MFG. R | 7. 11 | 3. 42E-11 |  | SFGdor. L | -<br>68. 23 | 0 |
|  | SFGdor. R | 4. 948 | 1. 11E-05 |  | SFGdor. R | -<br>93. 95 | 0 |
| 12 | MFG. L | -5. 55 | 5. 50E-07 | 21 | IPL. R | 3. 885 | 2. 64E-04 |
|  | MFG. R | -4. 926 | 1. 23E-05 |  | MFG. L | -<br>4. 797 | 6. 94E-06 |
|  |  |  |  |  | MFG. R | 4. 554 | 2. 01E-05 |
|  |  |  |  |  | SFGdor. L | -<br>21. 78<br>9 | 4. 15E-98 |
|  |  |  |  |  | SFGdor. R | 7. 121 | 1. 20E-11 |

Abbreviations: IPL: Parietal Inf lobule; MFG: middle frontal gyrus; SFGdor: dorsal superior frontal gyrus; R: right; L: Left.

**Table 6.**
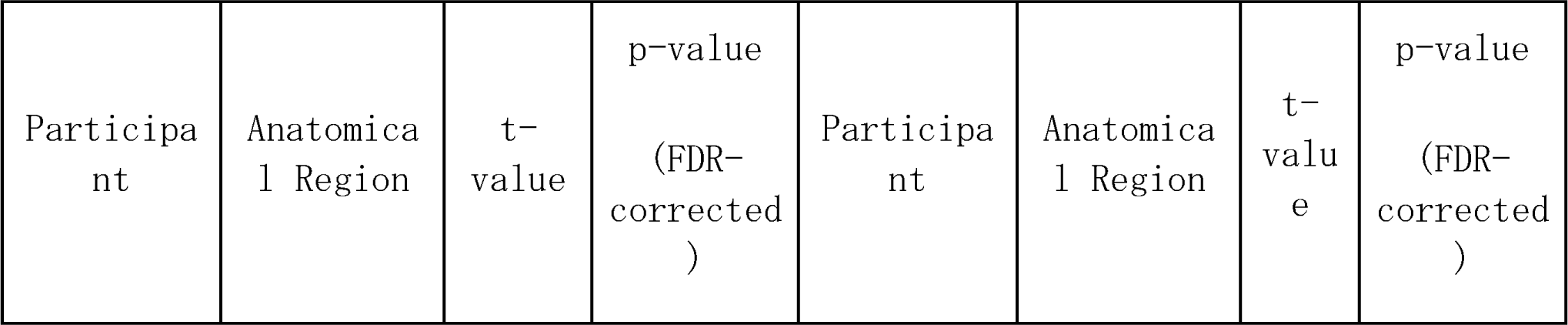
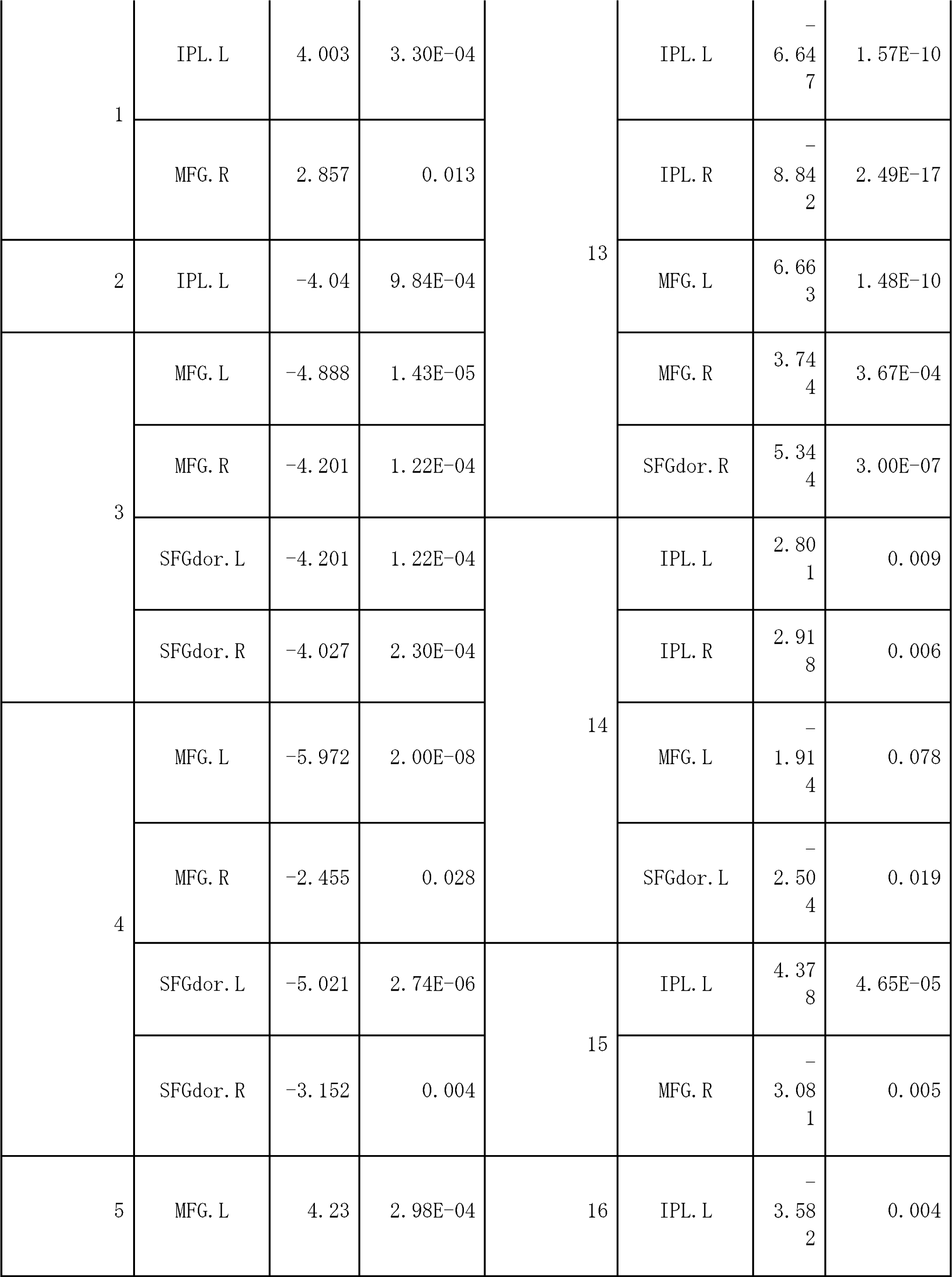
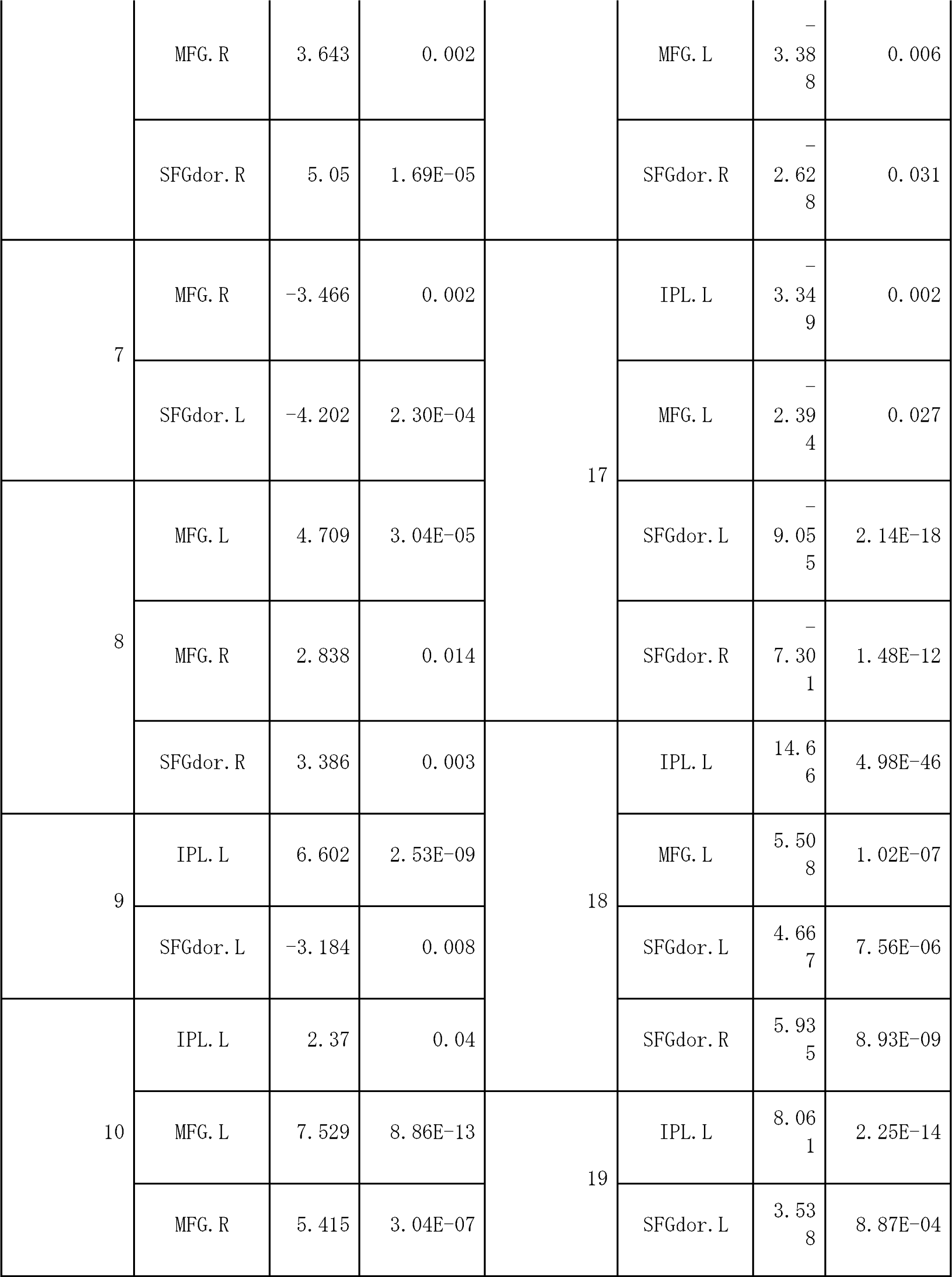
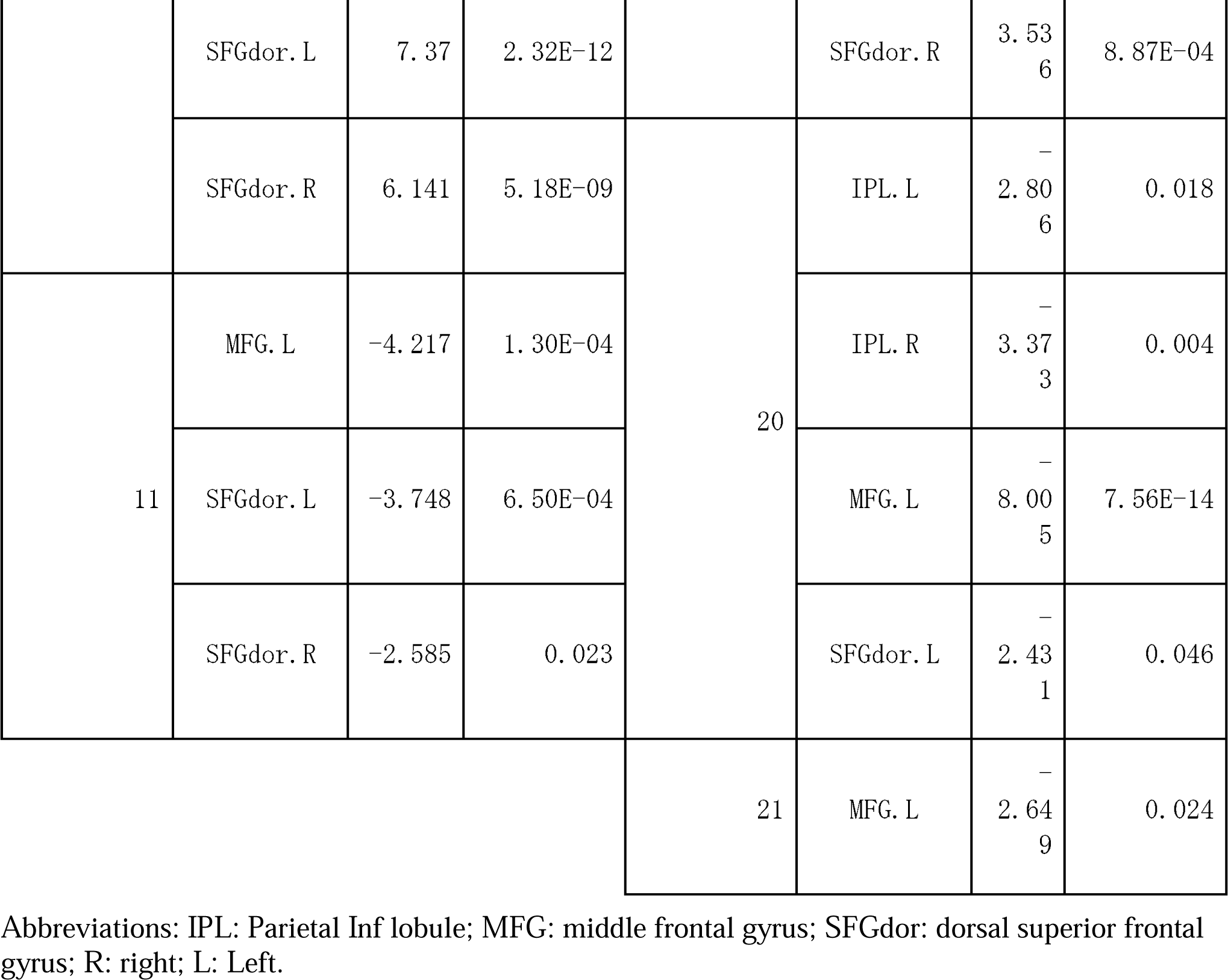
Brain regions that showed task-related activation for each individual in the planning block (non-expert older group). Only regions of interest with p < 0.05 are shown.

**Table 7.**
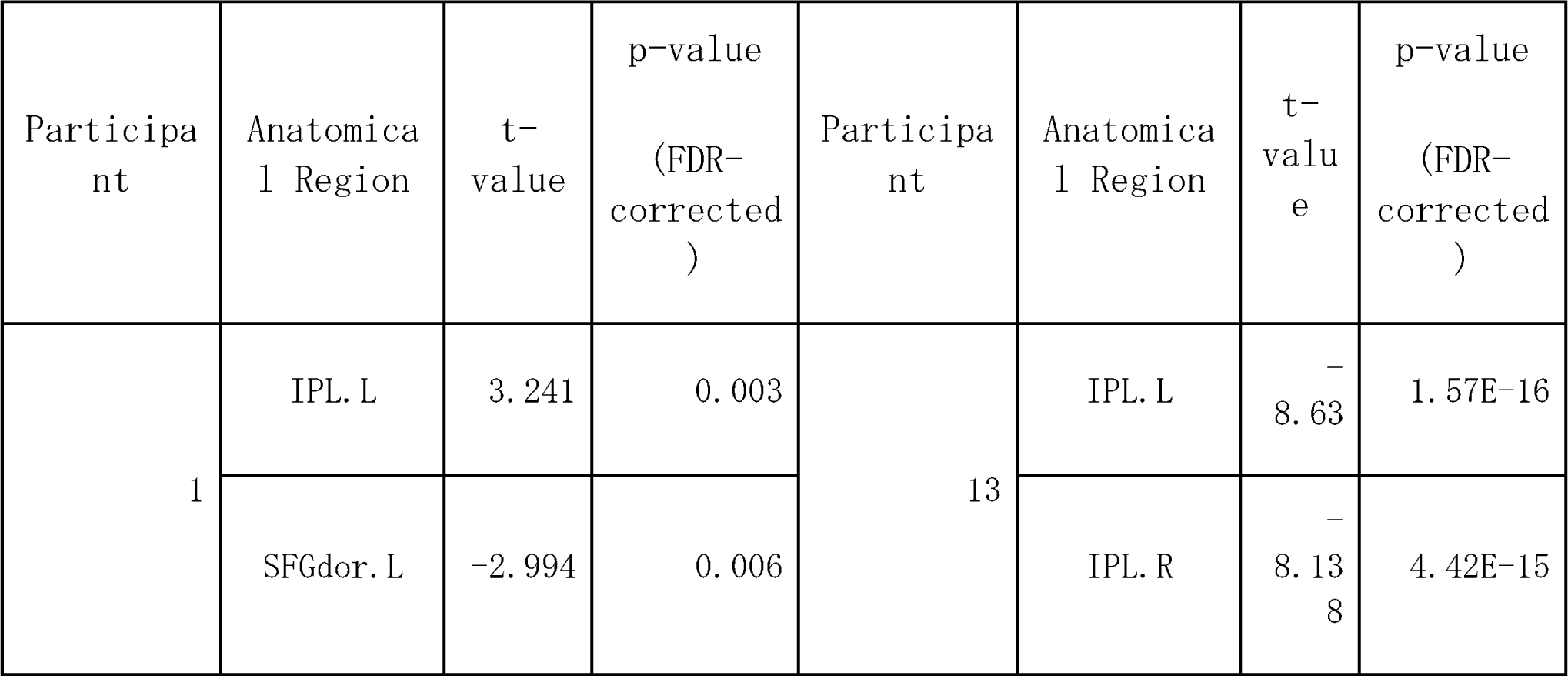
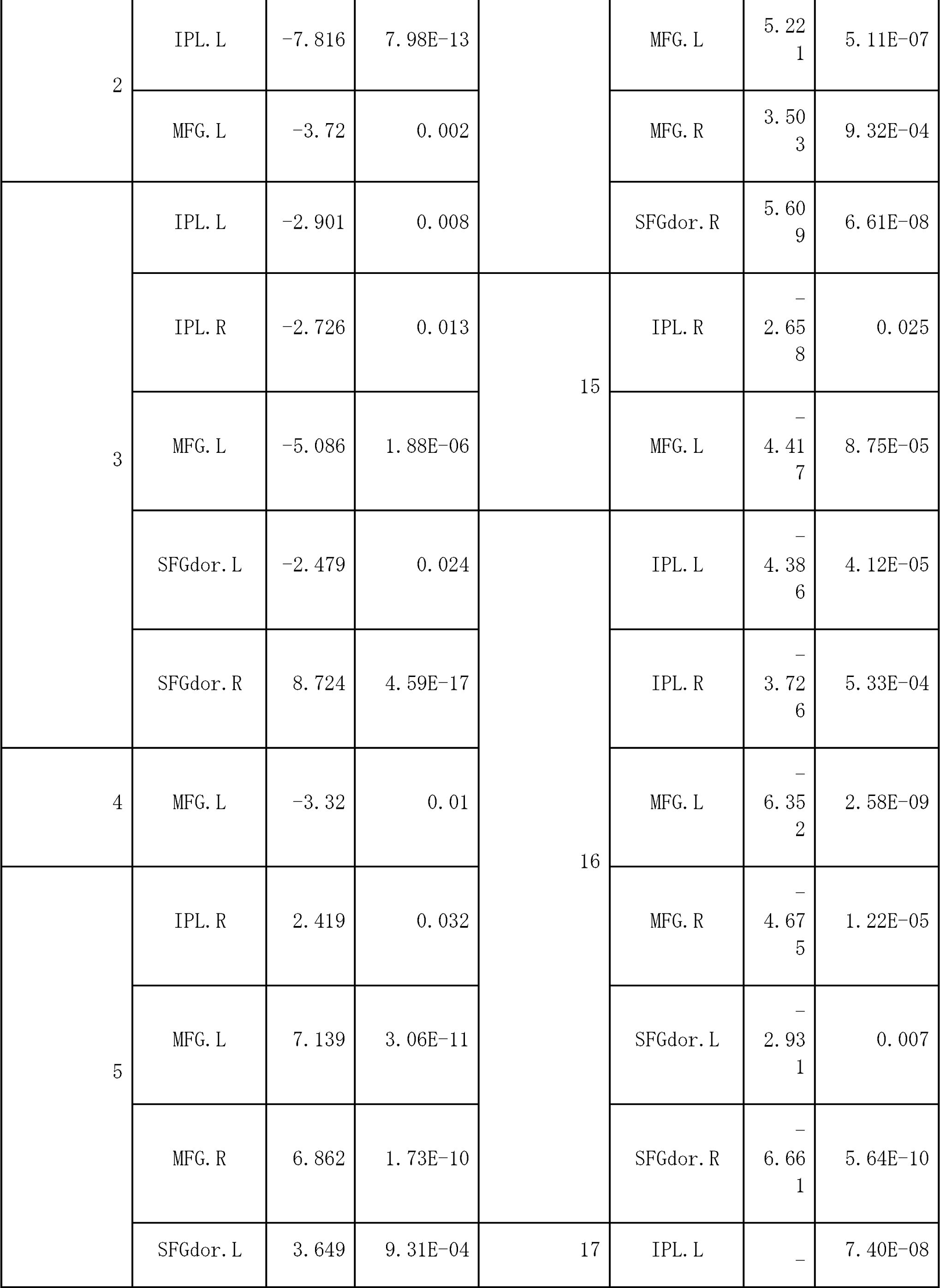
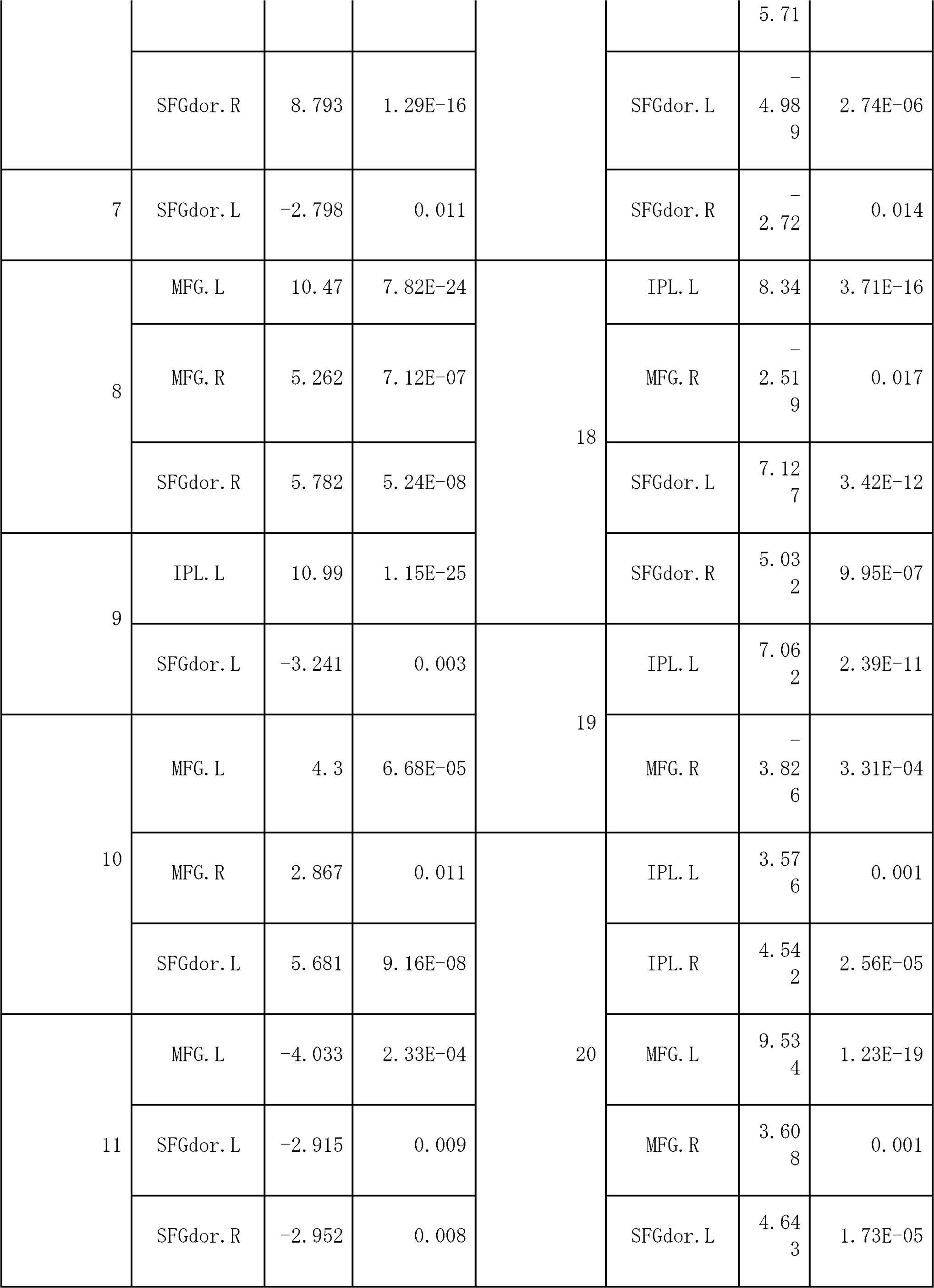
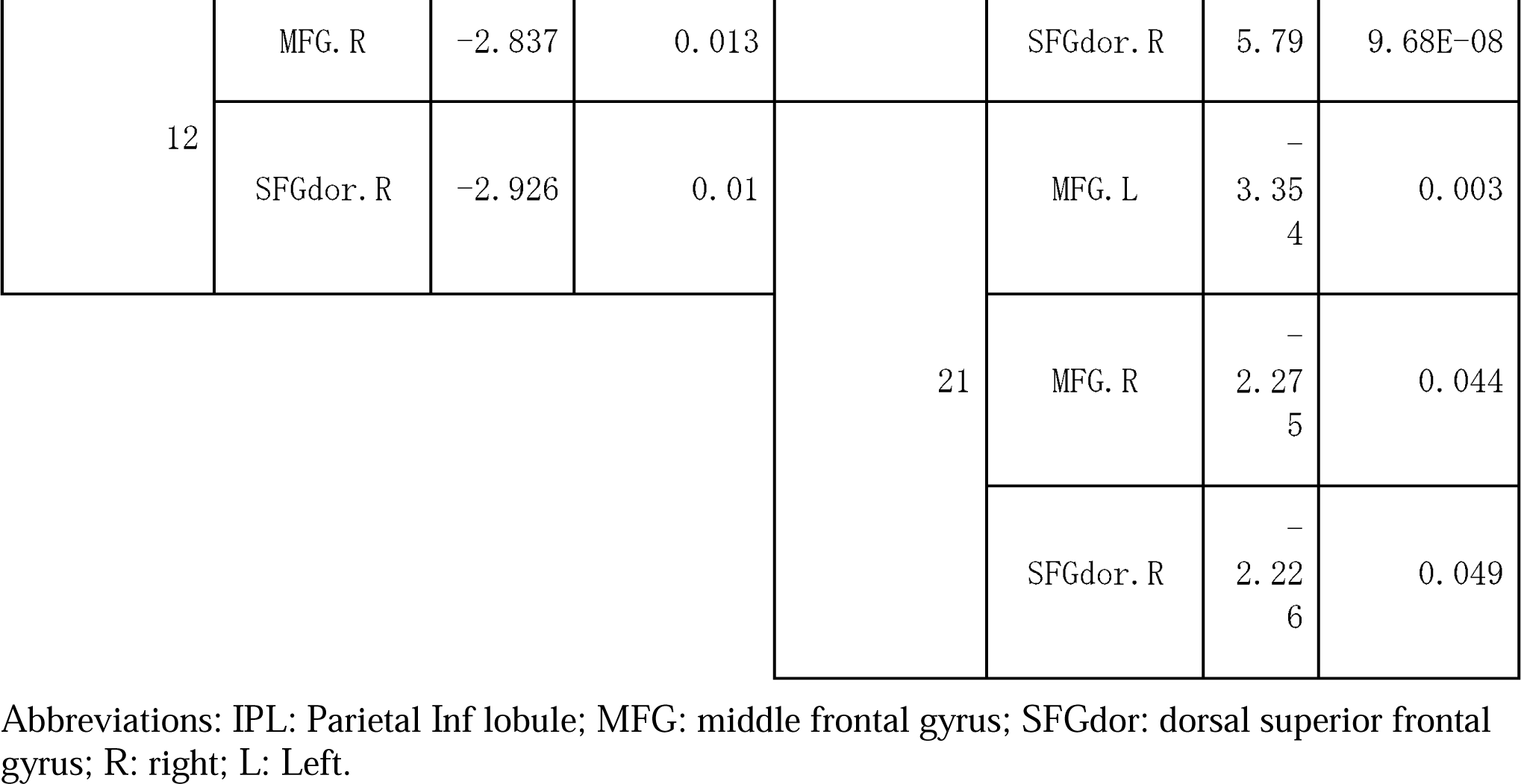
Brain regions that showed task-related activation for each individual in the calculation block (non-expert older group). Only regions of interest with p < 0.05 are shown.

**Table 8.**
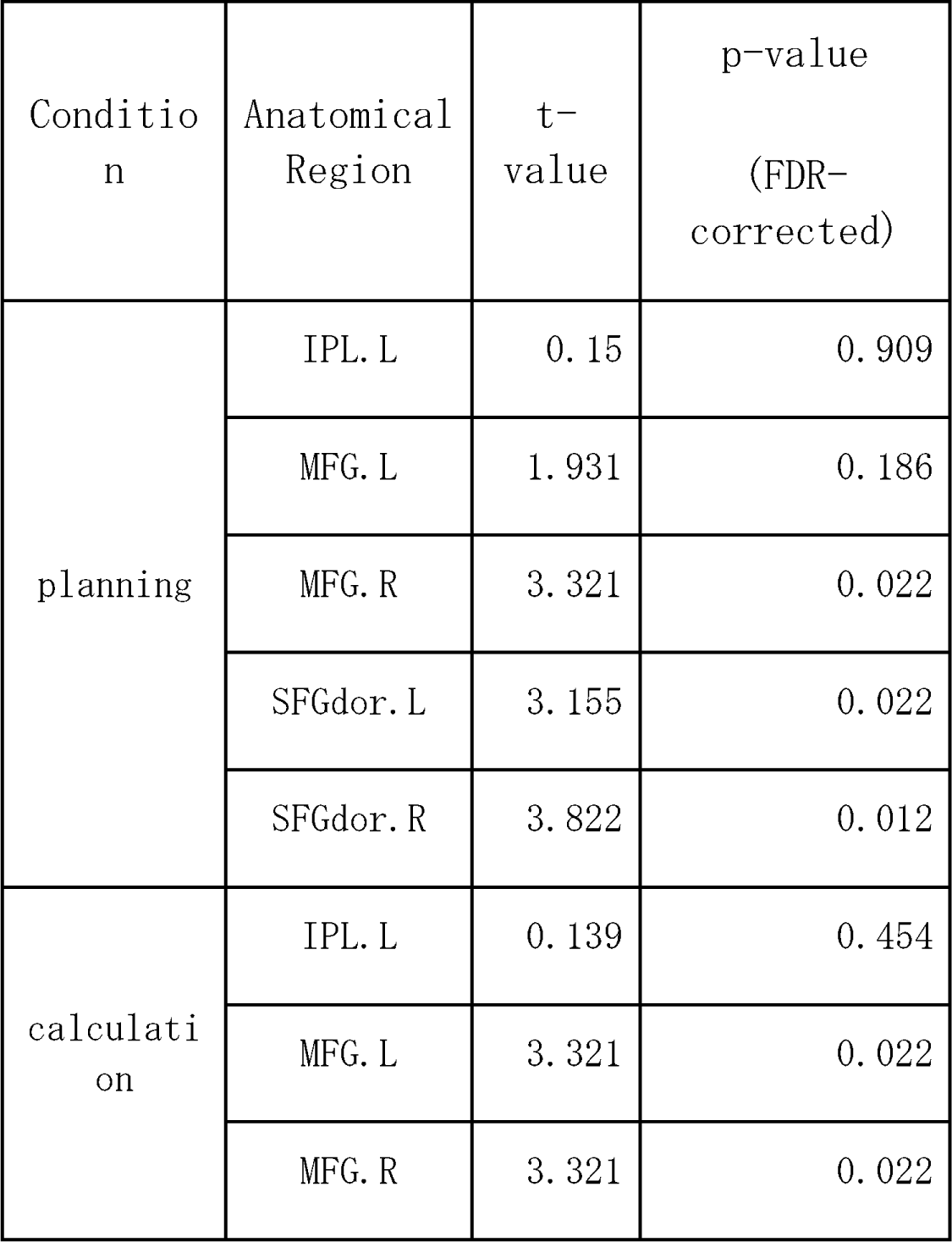
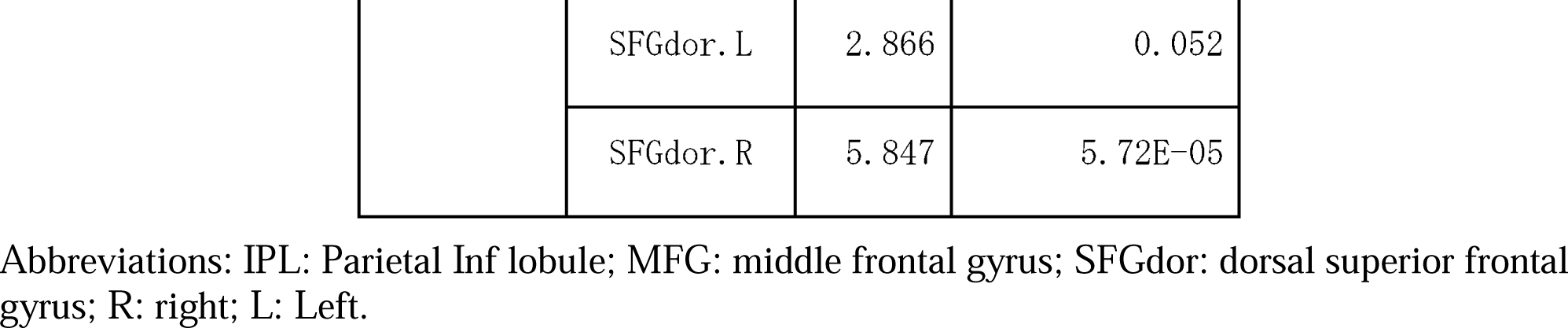
Task-related activation in the region of interest revealed by group-level analysis (younger group).

| Condition | Anatomical Region | t-value | p-value (FDR-corrected) |
| --- | --- | --- | --- |
| planning | IPL. L | 0.15 | 0.909 |
|  | MFG. L | 1.931 | 0.186 |
|  | MFG. R | 3.321 | 0.022 |
|  | SFGdor. L | 3.155 | 0.022 |
|  | SFGdor. R | 3.822 | 0.012 |
| calculation | IPL. L | 0.139 | 0.454 |
|  | MFG. L | 3.321 | 0.022 |
|  | MFG. R | 3.321 | 0.022 |
|  | SFGdor. L | 2. 866 | 0. 052 |
|  | SFGdor. R | 5. 847 | 5. 72E-05 |

Abbreviations: IPL: Parietal Inf lobule; MFG: middle frontal gyrus; SFGdor: dorsal superior frontal gyrus; R: right; L: Left.

**Table 9.** Task-related activation in the region of interest revealed by group-level analysis (expert older group).

| Condition | Anatomical Region | t-value | p-value<br>(FDR-corrected) |
| --- | --- | --- | --- |
| planning | IPL. L | 0. 127 | 0. 988 |
|  | IPL. R | 1. 706 | 0. 548 |
|  | MFG. L | -0. 163 | 0. 988 |
|  | MFG. R | 1. 238 | 0. 709 |
|  | SFGdor. L | 0. 343 | 0. 988 |
|  | SFGdor. R | 1. 04 | 0. 817 |
| calculation | IPL. L | 0. 545 | 0. 723 |
|  | IPL. R | 0. 367 | 0. 82 |
|  | MFG. L | 0. 999 | 0. 495 |
|  | MFG. R | 1. 542 | 0. 289 |
|  | SFGdor. L | 1. 115 | 0. 272 |
|  | SFGdor. R | 0. 989 | 0. 495 |

Abbreviations: IPL: Parietal Inf lobule; MFG: middle frontal gyrus; SFGdor: dorsal superior frontal gyrus; R: right; L: Left.

**Table 10.** Task-related activation in the region of interest revealed by group-level analysis (non-expert older group).

| Condition | Anatomical Region | t-value | p-value<br>(FDR-corrected) |
| --- | --- | --- | --- |
| planning | IPL. L | 2.282 | 0.385 |
|  | MFG. L | 0.446 | 0.943 |
|  | MFG. R | -0.928 | 0.794 |
|  | SFGdor. L | -0.331 | 0.943 |
|  | SFGdor. R | -0.684 | 0.904 |
| calculation | IPL. L | 1.234 | 0.525 |
|  | MFG. L | 0.309 | 0.876 |
|  | MFG. R | -1.118 | 0.54 |
|  | SFGdor. L | -0.496 | 0.803 |
|  | SFGdor. R | -0.469 | 0.815 |

Abbreviations: IPL: Parietal Inf lobule; MFG: middle frontal gyrus; SFGdor: dorsal superior frontal gyrus; R: right; L: Left.

### 3.3 Cognitive tests

#### 3.3.1 Computational ability test

The mean computational test scores for the younger group, the expert older group, and the non-expert older group were 24.0 ± 3.8, 20.5 ± 3.8, and 17.1 ± 5.5, respectively. As shown in Figure 9, the expert older group had higher scores on the computational skills test than the non-expert older group (*t* = 2.275, *p* = 0.029, *d* = 0.719).

**Figure 9.**
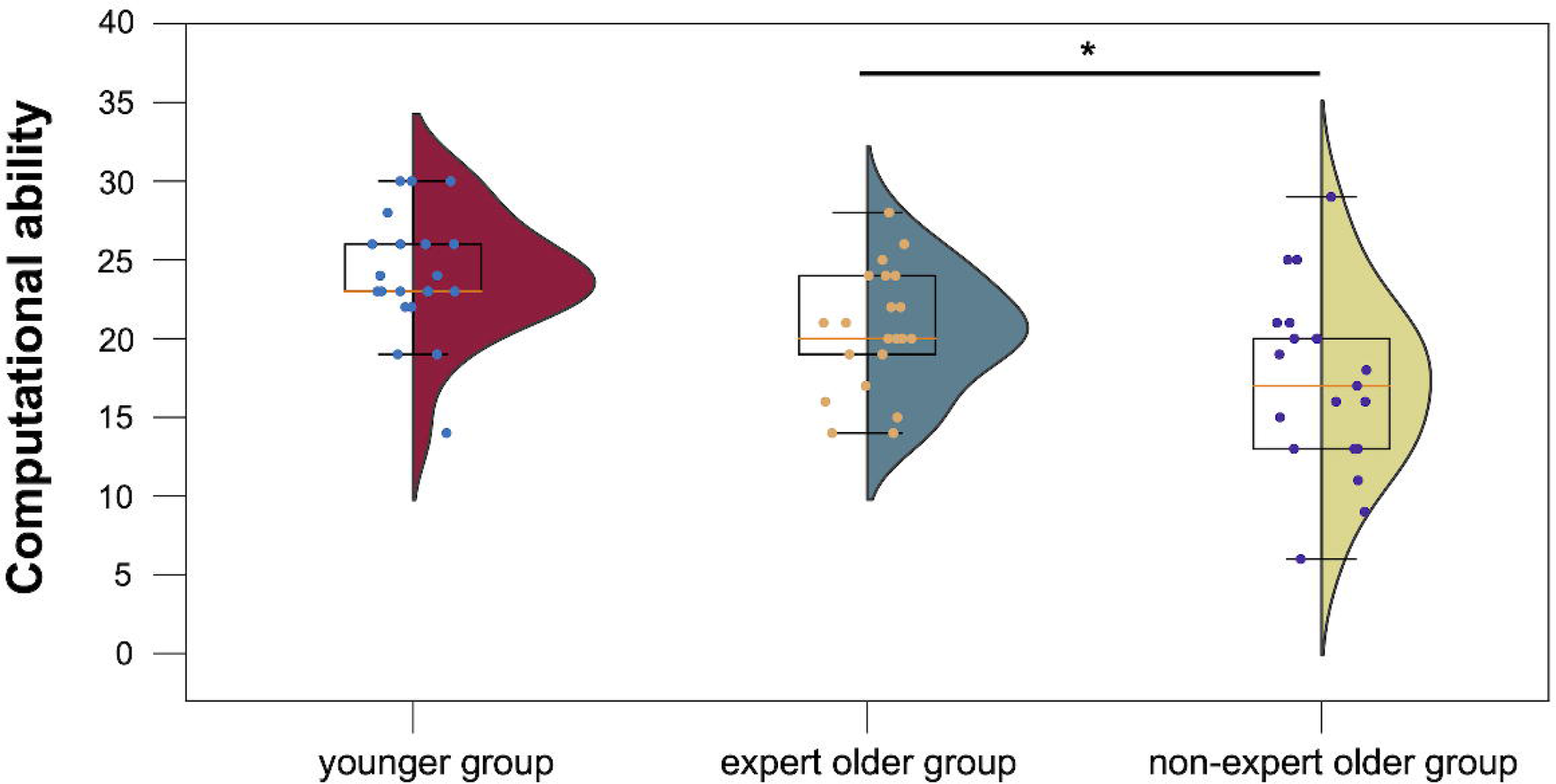
Computational ability test scores for the younger, expert older, and non-expert older group. *p <.05.

#### 3.3.2 STM test

The mean STM test scores of the younger group, the expert older group, and the non-expert older group were 74.6 ± 13.6, 57.1 ± 9.5, and 49.4 ± 9.9, respectively. As shown in Figure 10, the expert older group showed higher STM test scores than the non-expert older group (*t* = 2.475, *p* = 0.018, *d* = 0.792).

**Figure 10.**
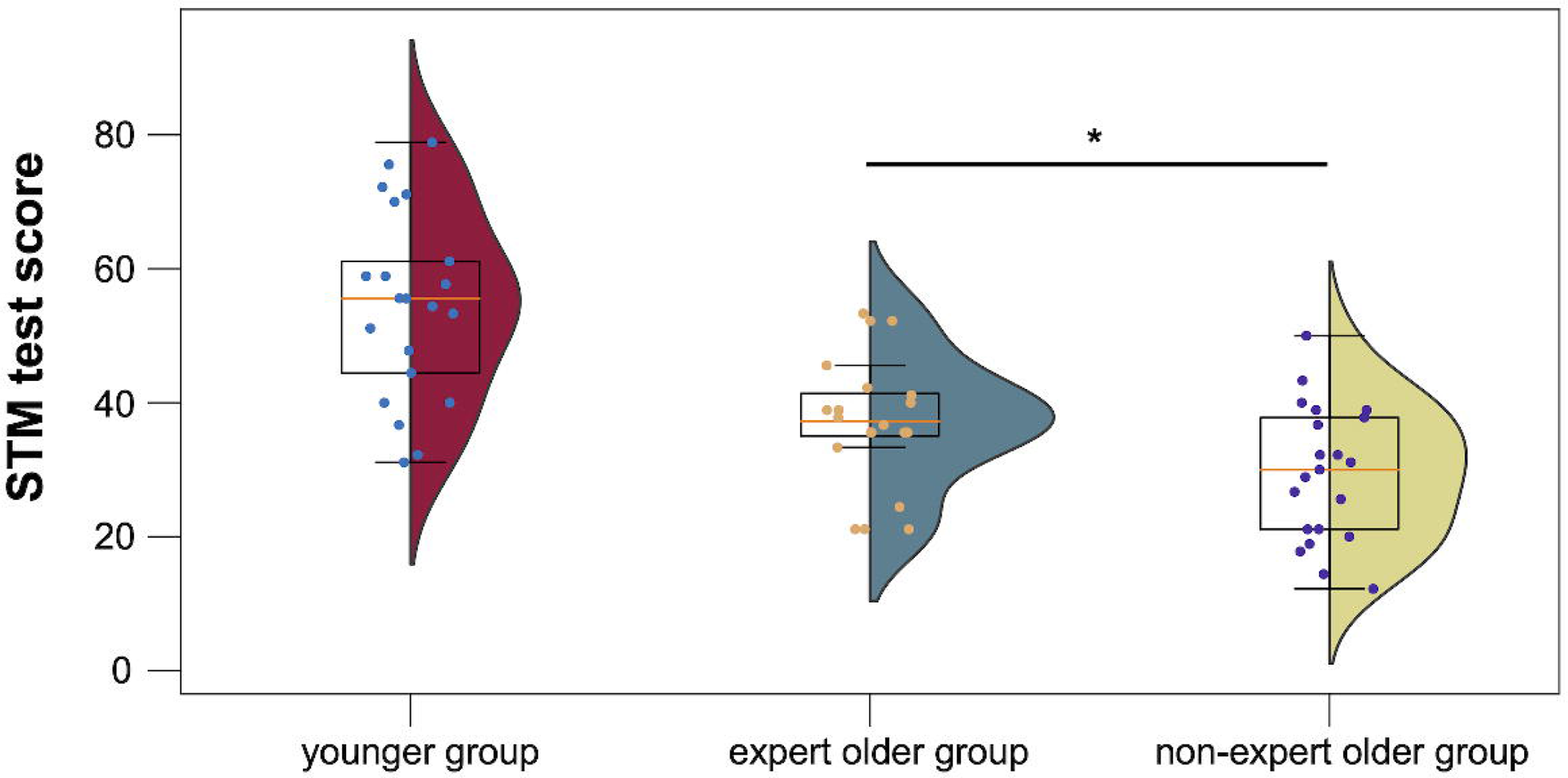
STM test scores for the younger, expert older, and non-expert older group. *p <.05.

#### 3.3.3 CogEvo

The scores of the five CogEvo tasks for each group of participants are summarized in Figure 11. In the expert older and non-expert older groups, there were no significant differences in the CogEvo scores (disorientation: *t* = 1.892, *p* = 0.067, *d* = 0.598, visual search: *t* = 1.435, *p* = 0.159, *d* = 0.454; flashlight: *t* = 1.386, *p* = 0.174, *d* = 0.438; number step: *t* = 1.722, *p* = 0.093, *d* = 0.550; just fit: *t* = 0.625, *p* = 0.534, *d* = 0.198).

**Figure 11.**
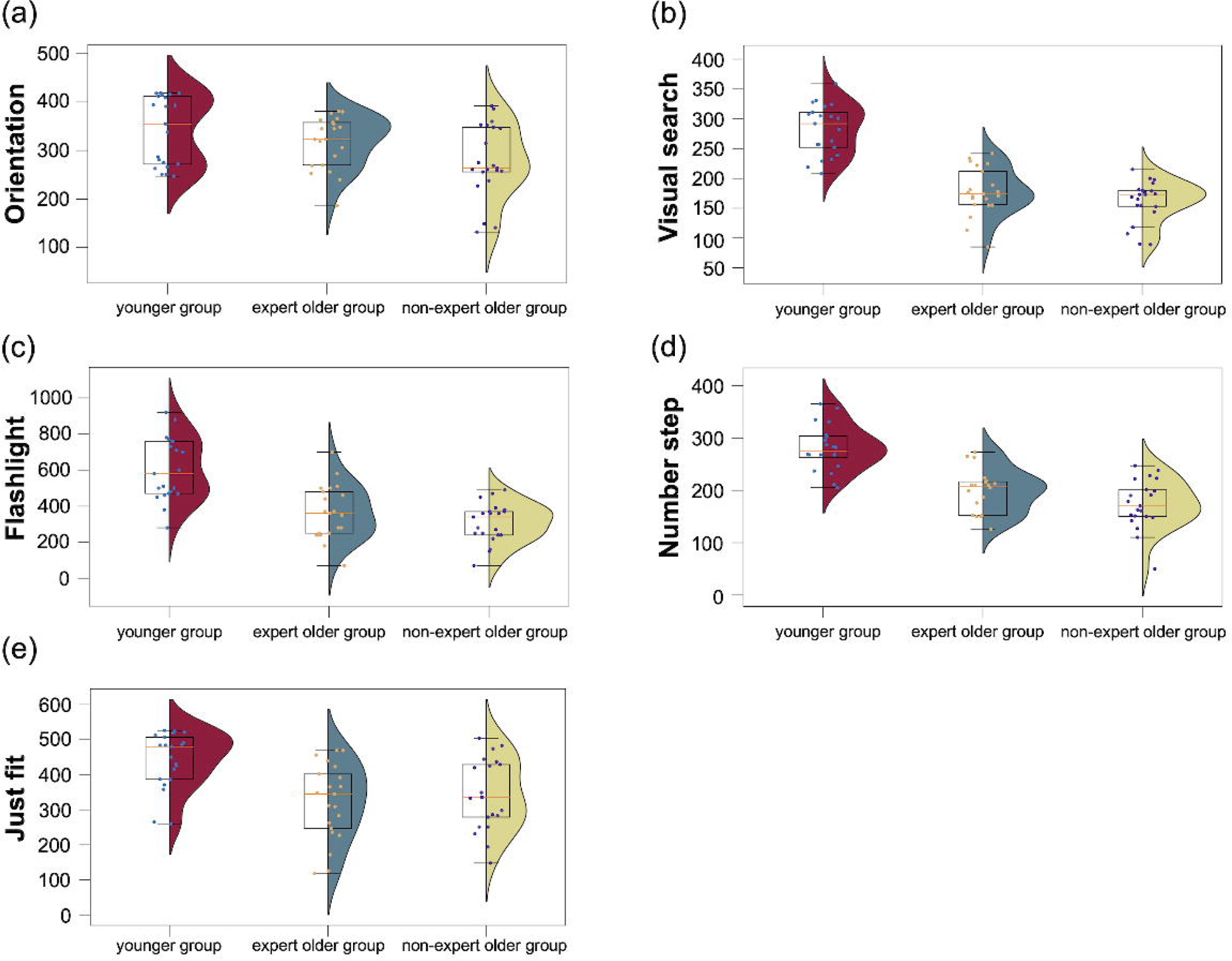
Scores on five different CogEvo tasks for the three groups of participants. (a) Orientation. (b) Visual search. (c) Flashlight. (d) Number step. (e) Just Fit.

#### 3.3.4 TMPT

The mean TMPT completion times for the younger, expert older, and non-expert older groups were ± 4.5, 58.1 ± 8.5, and 58.9 ± 9.2, respectively. There was no significant difference in TMPT completion time between the older and non-expert older groups (*t* = 0.272, *p* = 0.787, *d* = 0.086).

## 4 Discussion

### 4.1 Hypothesis 1

As seen in the results of the individual-level analysis in Tables 2-7, we were able to confirm the activation of SFGdor and MFG in the three participant groups. However, there was no significant activation of these regions in the group-level analysis. This may be because the mean activation level was estimated to be lower after averaging the results of the group as the individual activated areas varied even in the same region due to the distinct differences in the arrangement of measurement channels in the fNIRS device.

On the other hand, Figure 3 (d) shows that the LI values of the younger group and the expert older group were significantly higher than those of the non-expert older group in the planning block. In addition, there was no difference in LI values between the younger group and the expert older group. This indicates that the brain activation patterns of participants in the younger group and the expert older group are hemispherically lateralized during the planning of dart-throwing and while determining where to hit. The β-map in Figure 3 (a)-(c) also shows that the activation levels in the younger and expert older groups were higher in one hemisphere, while the non-expert older group showed no hemispheric difference. In the non-expert older group, there is no difference between the hemispheres, and bilateral activation levels of SFGdor and MFG are high. This result is consistent with a previous study in which left-right hemispherical asymmetry increased when cognitive training was performed in healthy older adults [12]. In summary, although SFGdor and MFG were not significantly activated in the expert older group in group-level analysis, Hypothesis 1 was established during the planning of dart throwing, suggesting that continual Wellness Dart training promotes hemispheric lateralization in brain activity in older adults.

### 4.2 Hypothesis 2

Figure 8 indicates that there were no significant differences in the β-values of the left and right IPL between the younger group and the expert older group. Further, there was no significant difference in the β-values of the left and right IPL between the younger and non-expert older groups, and between the expert and non-expert older groups. However, in the three participant groups in the present study, the IPL activation was comparable, suggesting that there was no difference in IPL activation between the older groups. The previous study to which the hypothesis refers involved healthy subjects and patients with MCI and Alzheimer’s disease [19]. The participants in the present study were only healthy adults, suggesting that the difference in cognitive function between the older groups and the younger group did not affect IPL activation.

### 4.3 Hypothesis 3

As shown in Figure 5, there was no positive correlation between the number of days of Wellness Dart experience and the LI value of the expert older group; therefore, hypothesis 3 was not supported. On the other hand, in Figure 6, the dart performance of the expert older group was higher than that of the other two groups. In addition, Figure 7 implies a positive correlation between dart performance and LI value in the planning block, although it is not statistically significant. The total score of our dart performance test indicates the accuracy of dart throwing, which is improved by the number of days of experience. Therefore, the results of Figures 6 and 7 may contribute to the validity of hypothesis 3. Also, since the current results suggested weak correlations, the correlations should be tested again with a larger sample size. These issues would be investigated in future longitudinal controlled studies of the Wellness Darts intervention that measure dart performance levels to verify the increase in LI values promoted by Wellness Dart training.

### 4.4 Cognitive tests

In addition to hypotheses testing, the results of three cognitive function tests to determine the impact of Wellness Darts on cognitive function are also discussed.

#### 4.4.1 Computational ability test

The score of the expert older group was significantly higher than that of the non-expert older group. In other words, the expert older group had better computational ability than the non-expert older group. According to previous studies, cognitive function, particularly for high-load tasks, declines as the brain deteriorates with age [40]. Therefore, the results of the computational ability test in this study suggest that continual practice of Wellness Darts can reduce the decline of calculation ability due to aging.

#### 4.4.2 STM test

The STM test scores of the expert older group were higher than that of the non-expert older group, and the difference was significant. Fandakova et al. [41] have shown that short-term memory capacity declines with age. Therefore, our results indicate that constant training with Wellness Darts can reduce the age-related decline in short-term memory capacity.

#### 4.4.3 CogEvo

No significant differences were found in the scores between the expert older group and the non-expert older group on the five CogEvo tasks. Although CogEvo has been reported to be able to significantly distinguish between Alzheimer’s disease, MCI, and cognitively normal groups of older adults [28, 29], our results showed that the expert older group and the non-expert older group did not differ significantly in their scores on the five CogEvo tasks. In fact, our results indicated that the expert older and non-expert older groups had similar cognitive levels. In other words, the older groups in our experiment had normal levels of cognitive function or the differences in cognitive function were at levels that were undetectable by CogEvo. Another explanation could be that the older groups were unfamiliar with the operation of CogEvo as it was conducted using a tablet device.

#### 4.4.4 TMPT

As seen in Figure 12, there was no significant difference in the completion time of TMPT between the expert older and the non-expert older groups. TMPT requires skill from the participant as a component of physical function. We expected that participants engaged in ongoing Wellness Darts exercises would demonstrate improved skillfulness; however, in this cross-sectional study, no differences were observed between the expert older and non-expert older groups. A previous longitudinal study that divided older adults into a dual-task (DT) exercise group and a control group observed a significant difference in TMPT before and after intervention in the DT exercise group, but no significant difference was identified in TMPT between the two groups [42]. Due to the pilot nature of the current study based on a cross-sectional comparison, a possible future direction could be to investigate whether the Wellness Darts intervention contributes to improving the TMPT completion time in longitudinal randomized controlled trials.

**Figure 12.**
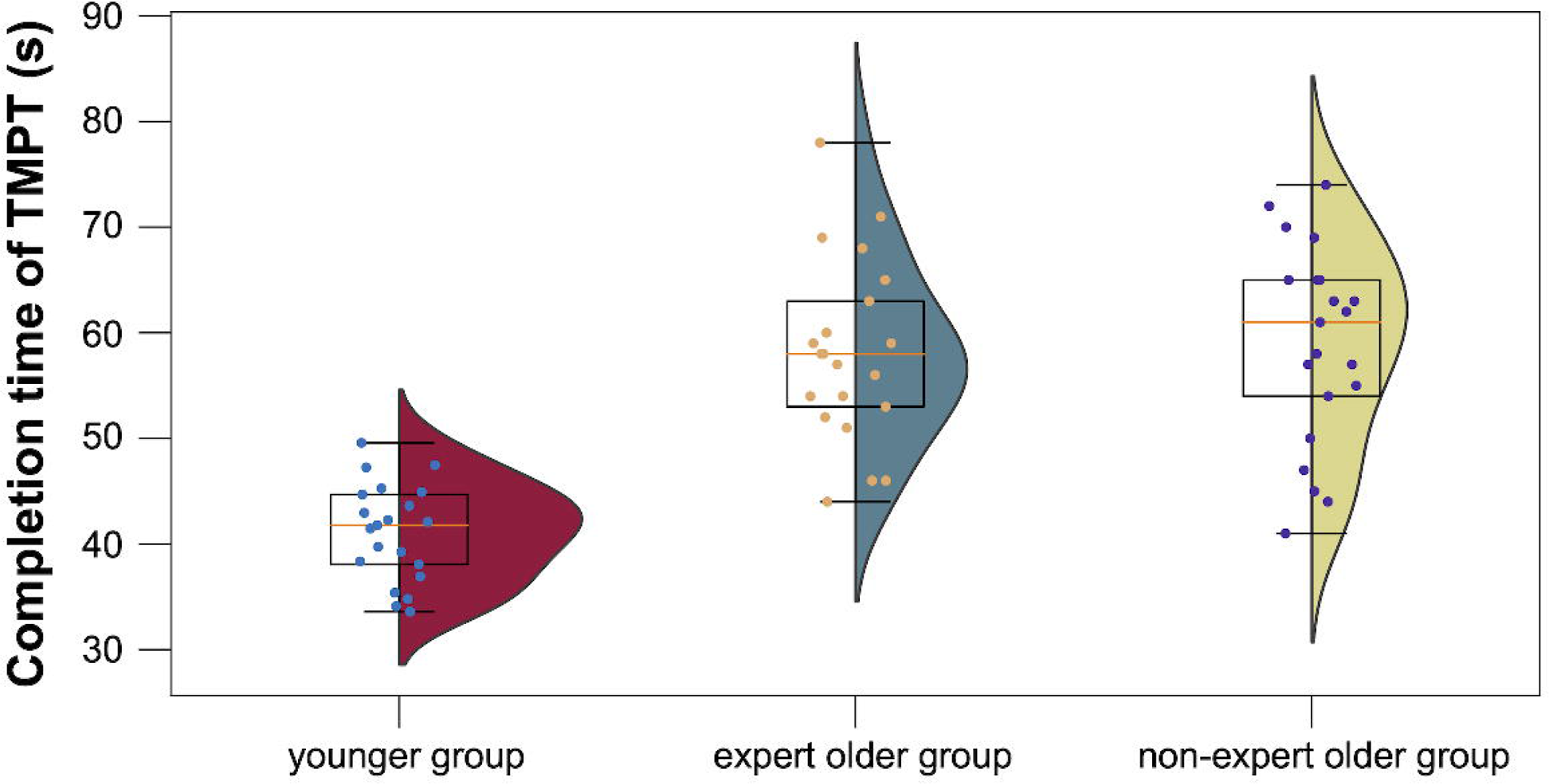
TMPT completion time for the younger, expert older, and non-expert older group.

### 4.5 Strengths of the current study, limitations, and future direction

Based on our findings, only hypothesis 1 was supported in the planning of dart throwing and in determining where to hit. Due to the pilot nature of this study, the sample size was limited, and the effects of several unseen parameters such as education level, socio-economic factors, body mass index, and smoking were not controlled and investigated. These limitations make the current study insufficient to demonstrate that Wellness Darts truly affected brain lateralization. However, despite these limitations, the current study revealed that older adults with Wellness Dart experience have a higher tendency toward hemispheric lateralization than those with no experience. The current results are valuable in motivating us to proceed to future longitudinal randomized controlled trials to explore the effects of continual Wellness Dart training on hemispheric lateralization with a larger sample size.

## 5 Conclusions

Wellness Darts is a cognitively challenging sport that combines exercise and calculation. Here we aimed to cross-sectionally confirm the difference in hemispheric lateralization between expert and non-expert players as a pilot study prior to the longitudinal randomized controlled trials. The results showed that the younger and the expert older groups had significantly higher LI values than the non-expert older group and that these values did not differ between the expert older and the younger groups. These results suggest that the Wellness Darts possibly promote hemispheric lateralization.

## 6 Declarations

### 6.1 Ethics approval and consent to participate

Studies involving human participants were reviewed and approved by the Research Ethics Committee of Doshisha University, Kyoto, Japan (approval code: 20014), and conducted in accordance with the Declaration of Helsinki. All participants were informed about the experimental method as well as the risks and signed written informed consent forms to participate in the study.

### 6.2 Consent for publication

he participants provided their written informed consent prior to enrolment in the study.

### 6.3 Availability of data and materials

The datasets generated during and/or analyzed during the current study are available from the corresponding author upon reasonable request.

### 6.6 Authors’ contributions

KoT, SH, TH, and MT contributed to the conception and design of the study. KoT and SH wrote the manuscript. KoT, SH, and MT conducted experiments. KoT performed data analysis. SH, KeT, TH, and MT advised on the data analysis and experimental design. All authors contributed to the manuscript revision and read and approved the submitted version.

## 6.7 Acknowledgments

We are deeply grateful to the members of the Kyotanabe Doshisha Sports Club (KDSC) for their cooperation in the experiment. We would like to thank Editage (www.editage.com) for English language editing.

